# Temporally Resolved and Interpretable Machine Learning Model of GPCR conformational transition

**DOI:** 10.1101/2025.03.17.643765

**Authors:** Babgen Manookian, Elizaveta Mukhaleva, Grigoriy Gogoshin, Supriyo Bhattacharya, Sivaraj Sivaramakrishnan, Nagarajan Vaidehi, Andrei S. Rodin, Sergio Branciamore

## Abstract

Identifying target-specific drugs remains a challenge in pharmacology, especially for highly homologous proteins such as dopamine receptors D_2_R and D_3_R. Differences in target-specific cryptic druggable sites for such receptors arise from the distinct conformational ensembles underlying their dynamic behavior. While Molecular Dynamics (MD) simulations has emerged as a powerful tool for dissecting protein dynamics, the sheer volume of MD data requires scalable and unbiased data analysis strategies to pinpoint residue communities regulating conformational state ensembles. We have developed the Dynamically Resolved Universal Model for BayEsiAn network Tracking (DRUMBEAT) interpretable machine learning algorithm and validated it by identifying residue communities that enable the deactivation of the β_2_-adrenergic receptor. Further, upon analyzing dopamine receptor dynamics we identified distinct and non-conserved residue communities around the contacts F170^4.62^_F172^ECL2^ and S146^4.38^_G141^34.56^ that are specific to D_3_R conformational transitions compared to D_2_R. This information can be tapped to design subtype-specific drugs for neuropsychiatric and substance use disorders.

## Introduction

G-protein coupled receptors (GPCRs) represent the largest membrane bound receptor family in the human genome, consisting of approximately 800 members.(*1*) GPCRs are expressed throughout the body and play crucial roles in signal transduction by responding to a wide range of external signals, including photons, ions, neurotransmitters, hormones, and proteins.(*2*) Due to their involvement in numerous physiological processes, GPCRs are linked to various diseases and nearly 30% of drugs on the market target this receptor family.(*2*) A majority of such drugs are small molecules or peptides that bind to the orthosteric site that is conserved in sequence and structure. Prominent drug target receptors, including for example, incretin receptors (GLP1s), dopaminergic, muscarinic, and serotonergic receptors, have functionally distinct isoforms with a high degree of sequence similarity in the orthosteric site. The non-conserved structural regions of highly homologous receptors are often located in disordered or unstructured loop regions that are dynamic. The functional relevance of the disordered regions often emerges from their dynamic ensemble of states. Hence, the realizable therapeutic efficacy of many GPCR therapeutics directed at specific isoforms is limited due to lack of target specificity. To address this challenge, we investigate isoform-specific information embedded in the structural dynamics of GPCRs.

GPCRs are dynamic proteins and in response to agonist and trimeric G-protein binding they undergo structural transition from inactive to active states, leading to activation of the G-protein triggering downstream signaling. While the distinct conformational states of GPCRs, such as inactive, active and intermediate states have been identified using high resolution cryo-electron microscopy (cryo-EM) and X-ray crystallography,(*3*, *4*) characterizing the transitions among these states at an atomic level using NMR, single molecule FRET, HDX and DEER is challenging due to the lack of atomic level spatial and temporal resolution of these measurements for membrane proteins. Hence, very often the results from these experimental techniques are integrated with Molecular Dynamics (MD) simulations to study GPCR activation and deactivation processes at an atomic level.(*5*) Valuable insights have been gained on residues that undergo conformational changes between active and inactive state ensembles in several GPCRs using MD simulations and their analyses.(*6–10*)

Since long time scale MD simulations are accessible, MD simulations that recapitulate large scale state transitions have become feasible. The strength of MD simulations lies in its ability to provide detailed atomistic level information on the ensemble of states and the transitions between them. While MD has emerged as a complementary tool to dissect GPCR dynamics, the sheer volume of data in multi-scale molecular dynamics simulations requires novel scalable and unbiased strategies to reveal structural insights into residue communities that regulate conformational transitions.

The recent wave of artificial intelligence (AI) and machine learning/deep learning (ML/DL) has led to great progress in novel applications to protein structure prediction(*11*) and dynamics.(*12*, *13*) Developments in ML/DL, including neural networks(*14*) and more sophisticated forms of non-deep learning predictors,(*15*, *16*) have provided new toolkits for the task of investigating and predicting protein dynamics. Generative AI-based methods are being currently developed for enhanced sampling of protein conformational space.(*17*, *18*) Despite the predictive potential of these ML methods, their lack of interpretability is a limiting factor in applying them for gaining granular mechanistic insights into protein dynamics. Additionally, there is no time-resolved ML model that can be used to analyze the MD simulation trajectories to identify the residues that enable conformational transitions in proteins. In general, there is a lack of interpretable, fully data-driven unsupervised time-resolved ML methods that unveil patterns in the protein dynamics data at the atomistic scale.

Bayesian network model or BNM (a prominent class of probabilistic graphical models) is an interpretable unsupervised machine learning model that has proven to be effective in identifying even weak probabilistic relations within noisy data. BNM has been successfully applied to analyze diverse biomedical datasets.(*19–23*) Recently, we have successfully applied BNM in analyzing MD simulation trajectories of GPCR: G-protein complex dynamics,(*24*) and identified cooperative residue contacts in the GPCR:G protein interface that facilitate selective G-protein coupling.(*24*) However, currently there is no data-driven interpretable time-resolved ML approach to analyze time-dependent transition events embedded in MD trajectories that recapitulate the macro-state transitions. Building upon this BNM framework, we have developed an innovative method to map the probabilistic relations among GPCR contacts as a function of time. Here we introduce Dynamically Resolved Universal Model for BayEsiAn network Tracking (DRUMBEAT) which enables us to identify key residue communities with coordinated movements that enable and effect macro-state transitions, such as GPCR deactivation. In this study, we first demonstrate the ability of DRUMBEAT to recapitulate previously known experimental and evolutionary insights into the deactivation process of GPCR structural prototype, the β_2_-adrenergic receptor (β_2_AR). In a fully data-driven fashion, we inferred that the community of residues centered around the contact between C327^7.54^ and F332^8.50^ enables the deactivation transition to the inactive state and the transition is completed by the ionic lock contact between transmembrane helices 3 and 6 (TM3-TM6) R131^3.50^ and E268^6.30^. While these important transition enabler and effector residue contacts are highly conserved among paralogs of β_2_AR, they are not strictly conserved among adrenergic receptor family subtypes. Our study of deactivation mechanisms of the two highly homologous D_2_ and D_3_ dopamine receptors (D_2_R and D_3_R) showed that two residue communities in the extracellular and intracellular region that are involved in the D_3_R deactivation mechanism are D_3_R specific and not involved in the D_2_R deactivation. This proves that DRUMBEAT is useful for identifying target-specific residue communities that can be harnessed for isoform-specific modulator design.

## Results

The input to DRUMBEAT is properties extracted from MD simulation trajectories that show macro-state transitions. For the β_2_AR system, we obtained 14 MD trajectories of deactivation starting from the active state structure, each 2μs to 10μs long, from DE Shaw and associates.(*25*) For the dopamine receptors D_2_R and D_3_R, we performed all-atom MD simulations starting from the agonist and G protein bound active state structure of D_2_R (PDB: 8IRS)(*26*) and D_3_R (PDB ID: 8IRT)(*26*). We removed the agonists and the G proteins and performed 31 runs of MD simulations each extending to 1.1-1.4µs for D_3_R and 35 runs extending to 300-600ns for D_2_R. The simulation times are different for D_2_R and D_3_R since the deactivation transitions occurred at different times. The MD simulations were performed in POPC lipid bilayer at constant temperature (300K) and volume using GROMACS2022.1 and CHARMMff v36m as detailed in the Methods section. To ascertain if the trajectories show transitions from active to inactive states, we calculated the inter-residue distances that are the hallmark of activation in class A GPCRs. The distance between residues 3.50-6.34 and 5.58-7.53 (using GPCR data base residue numbering system) were calculated for each trajectory in every GPCR system and plotted as 2D density plots shown in Fig. S1 and S2. Trajectories 1-11 for D_2_R and 1-12 for D_3_R did show a transition from active to inactive state as indicated by the decrease in the distance between the residues 3.50 and 6.34 on transmembrane helices 3 and 6 (TM3 and TM6) and simultaneous increase in the distance between 5.58 and 7.53 in TM5-TM7.

### Dynamically Resolved Universal Model for BayEsiAn network Tracking (DRUMBEAT) applied to analyze temporal transition events in proteins

#### The DRUMBEAT Method

The details of DRUMBEAT are given in the Supporting Information. The DRUMBEAT method developed in this work consists of three steps as described in the schema in Figure 1. The first step is to extract protein dynamics data such as pairwise residue contact information from MD simulation trajectories as the input for a machine learning model (Fig. 1A). Using the MD-specific BNM software BanDyT(*27*) we generate a BNM universal graph that yields the probabilistic dependencies between the input variables (residue contacts in this study). In this universal graph model, the residue contacts are the nodes and the edges are direct, non-transitive dependencies between pairs of contacts in the probabilistic space (Figure 1B). Finally, as shown in Fig. 1C, a DRUMBEAT model is contiguously re-computed along each trajectory using mutual information, and the high-ranking weighted degree measures are used to pinpoint critical events associated with state transitions in the protein. In principle, the method reported herein can be applied to any temporal or dynamic protein data and the first step may be augmented with different modalities depending on the nature of the available input data.

**Fig. 1.**
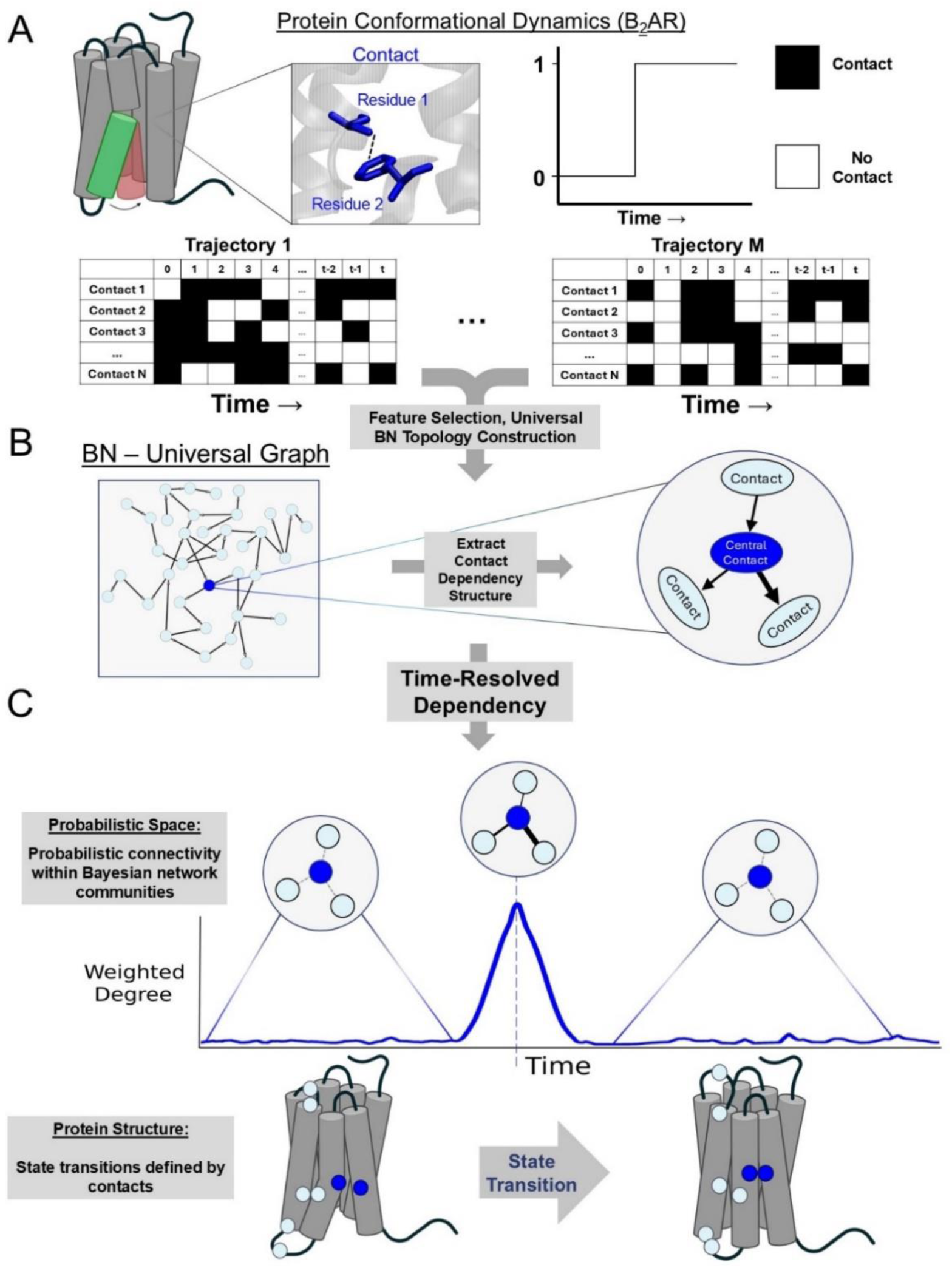
Workflow schema of DRUMBEAT. (**A**) Protein dynamics data serves as an input for the analysis pipeline. MD simulations data was converted to a flat binary residue contact dataset using the GetContacts algorithm. (**B**) All trajectories were aggregated, with representative sampling and filter-based feature selection, to create a universal graph via Bayesian network construction. The resulting graph is an interpretable probabilistic model that covers the transition of β_2_AR from active to inactive state. Embedded feature selection resulted in the identification of key probabilistic communities of contacts for further analysis. (**C**) A time-resolved Bayesian network was used to analyze transitions between protein conformations along the trajectories by continuously evaluating weighted degrees associated with key communities.

To build the BNM universal graph, we collectively analyze the aggregated 14 β_2_AR deactivation trajectories starting from an active state.(*25*) Each snapshot from MD simulations is expressed as a set of inter-residue contacts, assigning a value of 1 if the contact is formed, and 0 otherwise (Figure 1A). The choice of contacts as features is motivated by the fact that structural transitions in GPCRs, such as activation and deactivation, are actuated by cooperative changes in internal residue contacts. Analyzing co-dependencies among residue contacts will thus enable the understanding of allosteric modulation mechanisms across distant regions.(*28–32*) Using the matrices of contacts from multiple trajectories, we generate a BNM universal graph that comprehensively shows the significant direct local and non-local dependencies between residue contacts across the ensemble of trajectories.

Nodes in this BNM represent residue contacts, and the edges reflect the direct, non-transitive dependencies between pairs of contacts in the probabilistic space (Figure 1B). Some nodes show more connectivity over others, approximating a scale-free node connectivity distribution – not uncommon with biological BNMs(*33*) (Supplementary Figure S3). We quantified the connectivity of a node by its weighted degree (*D*_*i*_) which is the sum of all edge strengths connecting to a node *i*. Nodes with high weighted degree reflect contacts with the highest impact on the GPCR dynamics due to their probabilistic relations with many other contacts. In our previous work, we showed that nodes with high weighted degree represent cooperative contacts that allosterically regulate G-protein coupling(*24*).

Although the BNM universal graph is built from temporal data, the model does not explicitly have a temporal component. In other words, the connectivity in the network and corresponding strengths are a superset of all possible direct dependencies across the entire corpus of data (hence “universal graph”). We therefore incorporate a time-resolved weighted degree, *D*_*i*_(*t*), which measures the changes in probabilistic dependencies of individual contacts over time (Fig. 1C). To identify residue pairs that enable and effect the transitions we consider pairs of contacts given by the edges in the universal graph and continuously compute their dependency over time *via* mutual information in a time window that scans across each trajectory (more details in Methods). Each contact in the network is associated with an immediate probabilistic neighborhood, a **T**ime-**R**esolved **A**llosteric **C**ommunity or TRAC, where the strength of connectivity is temporally resolved and is given by *D*_*i*_(*t*) (Fig. 1C). We find that *D*_*i*_(*t*) of individual TRACs vary over time with regions of the trajectory where *D*_*i*_(*t*) is zero or very small, suggesting little or no dependency between nodes in the TRAC. Conversely, there are specific times at which *D*_*i*_(*t*) peaks, suggesting a moment of synchronous and cooperative movement of the constituent contacts of the TRAC in question. It is important to note here that the peaks in *D*_*i*_(*t*) do not correspond to distinct protein conformational states but rather serve as a signal denoting a transition between two states. As we show in the following sections, these contacts can mediate and/or facilitate the transitions that occur during the deactivation of the receptor. In subsequent sections, we will analyze the behavior of these TRACs over time in multiple trajectories and identify in a fully data-driven manner a small subset of TRACs whose contacts and residues are explicitly important in receptor dynamics and conformational transitions.

#### Application of DRUMBEAT to deactivation of β_2_-adrenergic receptor

We utilized 14 MD simulation trajectories of β_2_AR deactivation (with varying lengths ranging from two to ten microseconds), all of which contained a transition from an active (nanobody bound active state structure of β2AR) state to the inactive state.(*25*) To derive the BNM universal graph, each trajectory was encoded as a set of binary streams of temporally varying 1500-1700 unique inter-residue contacts as given by the program GetContacts(*34*) (example contacts represented by black lines in Figure 2A). To achieve a computationally tractable dataset for BNM convergence, we omitted contacts between neighboring residues and those having low mutual information with other contacts, retaining around 1000 contacts per trajectory (Fig. 2A, details in Methods). We then concatenated the individual binary datasets and further reduced the number of frames using representative subsampling, leading to a final dataset with 14,000 frames and 590 contacts. The resulting BNM universal graph derived from this dataset (590 nodes and 1390 edges representing direct dependencies between contact pairs) comprehensively describes the dependency structure among contacts for all the trajectories (Figure 2B). We highlight two of the contacts in the BNM universal graph showing the highest weighted degrees (*D_i_*), R131^3.50^ _E268^6.30^ and C327^7.54^_F332^8.50^, colored in blue and purple, respectively. The contact R131^3.50^ _E268^6.30^ was already previously shown to be a critical "micro-switch" that reduces in distance upon deactivation.(*28*) This serves as initial evidence of the method’s ability to identify contacts important for the transition.

**Fig. 2.**
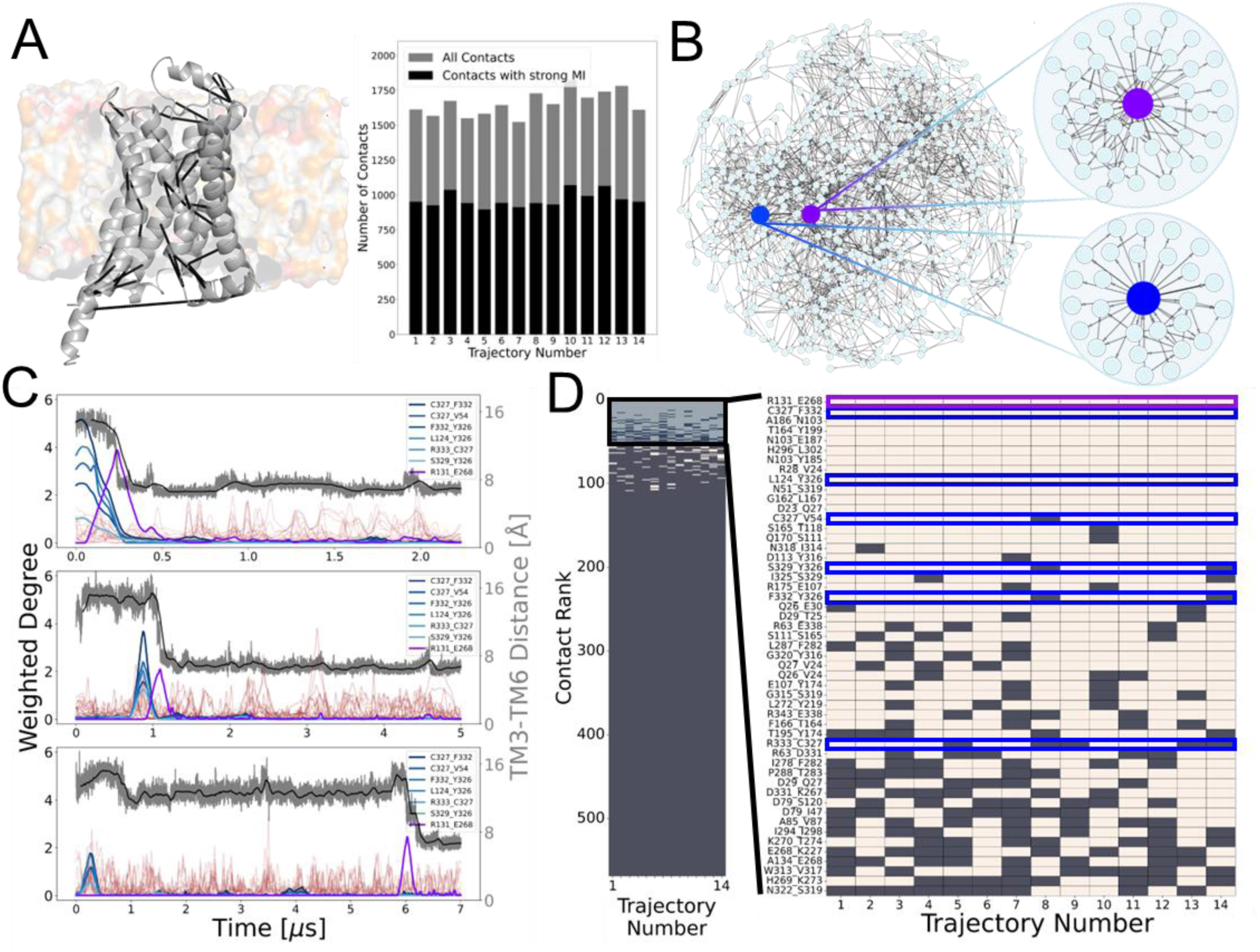
Construction of the universal graph from ensemble of trajectories, and subsequent time-resolved Bayesian network output. (**A**) Dynamics data in the form of MD trajectories of β_2_AR in lipid membrane transitioning from active to inactive state. Universal dataset built using contacts with strong mutual information with other contacts. (**B**) Universal graph output from the Bayesian network modeling, and corresponding time-resolved allosteric networks shown with the central node R131_E268 and C327_F332 highlighted in purple and blue, respectively. (**C**) Time-resolved Bayesian network output for trajectories 8,11 and 13 with main contacts highlighted in purple and blue. The remaining contacts are shown in brown. (D) Ranking of contacts by weighted degree above the threshold of 1 bit. The contacts shown in part C are highlighted, also in purple and blue.

### Analysis of the ensemble of β2AR transition trajectories reveals a subset of contacts as mediators of transition

To derive the time dependencies of each contact, we calculate its weighted degree *D*_*i*_, as a function of time for individual trajectories, using the BNM universal graph as a common topology (details in Methods). By monitoring the time evolution of weighted degree of each contact in individual trajectories, as described in the previous section, we explicitly determine *when* these probabilistic dependencies manifest during a trajectory and infer on the protein dynamics during transition.

For each separate trajectory the computed *D*_*i*_(*t*) for each node yields a series of peaks of varying size associated with the manifestation of the probabilistic neighborhood at various times throughout the trajectory. Figures 2C shows the results for three trajectories of β_2_AR (See Supporting Information Figure S5 for all 14 deactivation trajectories of β_2_AR). For each trajectory, *D*_*i*_(*t*) is plotted for the top 50 contacts ranked by amplitude. For reference indicative of the deactivation process in GPCRs, the interhelical distance between TM3 and TM6 helices is also plotted in grey. We identified a subset of the key contacts by applying three criteria: (1) signals with amplitude above one bit, or about one standard deviation, (2) occurrence of the same signal in all the trajectories, and (3) no more than one or two peaks in a single trajectory. The third criterion arises from our goal of analyzing macro-state transitions in the protein. Here, the two key signals remain (Figure 2C) --- the blue signal corresponds to a set of contacts with concentric peaks and the purple signal corresponds to a single contact and manifests at the transition defined by the drop in interhelical distance. The remaining signals, shown in brown, have less reproducible features across the trajectories and although they might contain potentially interesting and useful information on microstate transitions within the active or inactive state ensembles, they will not be the focus of this study.

To select the contacts in a fully data-driven manner, we ranked contacts based on the amplitude of weighted degree *D*_*i*_(*t*), as well as prevalence in multiple trajectories for all TRACs. Figure 2D shows the contact ranking in the form of a heat map of the amplitude of weighted degree *D*_*i*_(*t*), where the tan squares denote contacts with weighted degree above the threshold and brown squares otherwise. A large proportion of contacts failed to meet the threshold in even a single trajectory, reiterating the power law distribution for connectivity of contacts in this system. On the right, Figure 2D shows a zoom-in of the contact ranking for the top 50 contacts and highlights the contacts associated with the enabler and effector signals with the corresponding blue- and purple-colored boxes, respectively. The top-ranking contacts by these criteria are C327^7.54^_F332^8.50^ and R131^3.50^_E268^6.30^, and we will focus on these TRACs to identify the mechanisms by which the receptor deactivates.

When analyzing the signals in all the trajectories (Figures 2C and S5) we found that the contacts C327^7.54^_F332^8.50^ and R131^3.50^_E268^6.30^ (blue and purple signals, respectively) always appear and, interestingly, the peak of the C327^7.54^_F332^8.50^ contact always precedes the peak corresponding to R131^3.50^_E268^6.30^. Thus, we infer that the TRAC C327^7.54^_F332^8.50^ along with other concentric signals are the *enablers* and the TRAC R131^3.50^_E268^6.30^, the purple signal, is the *effector* of the transition from active to inactive states.

### The transition between conformational states in β2AR as seen via a change in residue community connected to enabler C327^7.54^ _ Y326^7.53^ and effector R131^3.50^_E268^6.30^ contacts

Figure 3A shows the probabilistic neighborhoods associated with the enabler C327^7.54^ _ Y326^7.53^ (blue) and effector R131^3.50^_E268^6.30^ (purple) TRACs. The concentric nature of the peaks that make up the enabler signal (Figure 3A; blue) is represented by a strongly interconnected probabilistic neighborhood with multiple overlapping residues. Specifically, the residues C327^7.54^ and Y326^7.53^ occur in multiple enabler contacts and have been shown by experiments to be important for the receptor function, with mutation in these residues resulting in increased or decreased agonist mediated β_2_AR activity.(*35–38*) Moreover, Y326^7.53^ is a part of the well characterized and conserved NPxxY motif that acts as a structural toggle, playing essential roles in rearranging the intracellular region for downstream signaling.(*28*, *29*, *32*, *39*, *40*) Second, and somewhat less obvious, is the closeness in probabilistic space between the two neighborhoods. There exists a set of 10 connector contacts that have an enabler and effector central node as an immediate neighbor. These connector contacts play a distinct and informative role in the conformational changes attributed to the enabler and effector signals as highlighted in this section. In addition, this degree of closeness in the probabilistic space between the enabler and effector TRACs (although distant in structural space) implies a strong probabilistic allosteric relationship between the two conformational changes that they represent. Interestingly, the closeness in probabilistic space does not definitively imply temporal or spatial closeness of the signals, as we highlight in an example below.

**Fig. 3.**
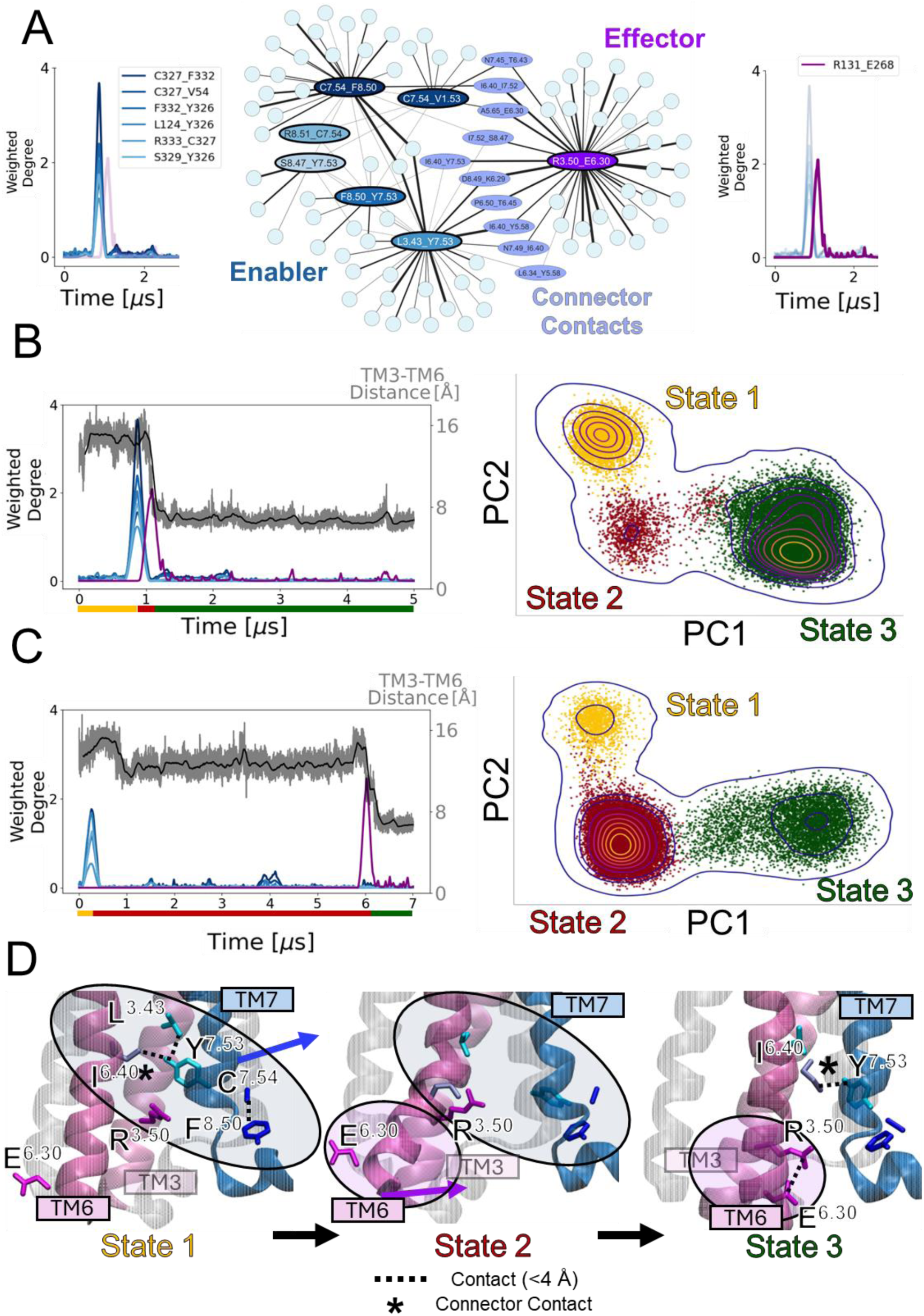
Analysis of “enabler” and “effector” signals shown in blue and purple, respectively. (**A**) Subnetwork corresponding to the enabler and effector TRACs with corresponding central nodes. The subnetwork shows a set of contacts that connect the two TRACs labeled as the “connector” nodes. (**B** & **C**) Conformational space analysis using two different trajectories and the contacts in the subnetwork. Principal component analysis reveals three distinct states in both trajectories. Coloring of the points in the PCA space with respect to the timepoints given by the peaks in the trajectory shows the blue peaks corresponding to the transition between State 1 and State 2, and the purple signal --- between State 2 and State 3. (**D**) Structural analysis of the three states taken from the trajectory in part **C**. Transition from State 1 to State 2 is characterized by the movement of TM7 away from TM3 resulting in breaking of contacts in the enabler TRAC. The subsequent transition from State 2 to State 3 is characterized by TM6 moving in towards TM3 resulting in the formation of the salt bridge between effector TRAC central contact residues R131^3.50^ and E268^6.30^.

As mentioned above, each TRAC signal represents the coordinated motion of a subsection of the protein structure defined by the constituent contacts in the respective TRAC. We declare these events of coordinated motion as transitions between distinct conformational states of the protein. Consider the trajectory shown in Figure 3B where there is an enabler signal and an effector signal. The presence of these two signals implies that two transitions are occurring in the trajectory between three distinct states. To further explore this idea, we carried out a principal component analysis (PCA) using the 98 contacts in the combined enabler and effector network in Figure 3A. Figures 3B and 3C depict the principal component analysis (PCA) results for two different trajectories and both show the presence of three states in the state space defined by principal component 1 (PC1) and principal component 2 (PC2). Moreover, the contacts with the highest weighting in PC1 overlap with the set of enabler contacts while the highest weighted contact for PC2 is the effector central contact. This suggests that the enabler signal represents the transition between state 1 and state 2 and the effector signal represents the transition from state 2 to state 3.

We can also analyze the temporal relationship between the two signals, and therefore the transitions. In principle, the population of individual states in the PCA space correlates with the time spent in that state as recapitulated by the MD simulations. The peaks provide a quantitative measure of the temporal adjacency of the state transitions, and therefore the states themselves. For instance, in Figure 3B, the enabler and effector signals are within one microsecond of each other. This ultimately corresponds to the low density of state 2 in the PCA analysis on the right. This temporal adjacency coincides with the closeness of the two communities in the network shown in Figure 3A. However, the trajectory in Figure 3C shows the enabler and effector signals nearly 6 μs apart, far away in time, and yet, its dynamics still corresponds to the closely connected probabilistic neighborhoods. This implies that even though the conformational changes in the protein occur at different time points, the enabler and effector residue communities remain the same. This is a key finding of this work.

Figure 3D shows the residue contacts involved in the enabler and effector signals in the three distinct states delineated from PCA. In state 1, the central contacts in the enabler TRAC break and make contacts involving residues C327^7.54^ and Y326^7.53^ as TM7 moves away from TM3. During this change, a large portion of breaking and forming contacts between C327^7.54^ and F332^8.50^ also occurs between TM7 and helix 8. This transition gives rise to state 2, an intermediate state between active and inactive conformations, a finding that follows the report(*25*) from which the trajectories were obtained. The precise structural motion and key residues involved provide a new and deeper understanding of the transitioning to the intermediate state. After some time in this intermediate state, the second transition associated with the effector signal occurs, leading to state 3. This transition occurs via TM6 swinging into the G-protein binding pocket towards TM3, ultimately forming a salt bridge between the central contact of the effector TRAC, R131^3.50^ and E268^6.30^, indicating the transition of receptor into inactive state.(*41*) Notably, the connector contact between residues I278^6.40^ and Y326^7.53^, initially broken during the first transition, is reformed in this state, reiterating its structural role in both transitions. Previous studies corroborate the importance of these key residues. For instance, the salt bridge between R131^3.50^ and E268^6.30^ is one of the established indicators of rhodopsin deactivation(*42*), and R131^3.50^ is part of conserved ED/RY motif on TM3 across all class A GPCRs.(*43*) Also, Y326^7.53^ contributes to GPCR activation via the conserved NPxxY motif.(*44*)

It is worth noting that the mechanistic conclusion for β_2_AR came from a fully data-driven unsupervised learning approach without any previous information or knowledge of the system (except for the interhelical distance, which serendipitously served as a ground-truth positive control). Taken together, our DRUMBEAT based fully data-driven analysis has uncovered the two transitions during deactivation and the precise residues involved in enabling these transitions.. We envision these types of findings manifesting from our methodology for any other *de novo* protein system where no prior information is available.

### Transition Enablers and Effector residues are conserved across adrenergic receptors

To investigate whether the residues identified as crucial during the deactivation transitions of β_2_AR are preserved in the adrenergic family, we have analyzed their conservation level in comparison to other residues, by computing the conservation score (C_i_) using Von Neumann entropy.(*45*) This method considers the chemical similarity between different amino acids to obtain a more reliable evaluation of the degree of conservation along the alignment. Briefly, it constructs the density matrix ρ from the relative frequencies of residues at each position and a similarity matrix (BLOSUM50). The Von Neumann entropy is *S*_*i*_ = − ∑_*i*_ λ_*i*_log_20_λ_*i*_ with λ_i_ the eigenvalues of ρ. Then, the conservation score is obtained by subtracting S_i_ from the maximum possible entropy (in our case 1), C_i_=1−S_i_. Using a multiple sequence alignment (MSA) of the 68 adrenergic receptor family sequences from 20 species from the GPCR database (https://gpcrdb.org/), we computed C_i_ for each position of β_2_AR in the alignment (Fig. 4A). As evident from Fig. 4A,B, key residues involved in the transition have higher conservation level compared to other receptor residues indicating their implication in β_2_AR stability and function. Numerous prior studies have demonstrated the importance of the residues identified by DRUMBEAT as enablers and effectors of transition. The residue R131^3.50^ is part of the conserved E/DRY motif and is responsible for modulating agonist binding and G protein coupling and governing receptor conformations.(*43*, *46*) In the CWxP motif, the residue P288^6.50^ is involved in the rotamer toggle switch mechanism that plays a role in active forms of GPCRs.(*47*, *48*) Moreover, the connector residues N322^7.49^ and Y326^7.53^ are part of the conserved NPxxY motif which contributes to GPCR activation and state transition.(*39*, *44*) The probability distribution function (PDF) of the conservation score in Figure 4D shows that the amino acid positions that are part of the enabler and effector central contacts(red), connector contacts that enable direct “communication” between those hubs in the probabilistic space(green), and the set of all the contacts in the first neighborhood of the main hub(yellow) are highly conserved compared to all the residues in the receptor. The rank-sum test shows a statistically higher level of conservation (higher values of C_i_) in DRUMBEAT-identified positions, indicating that evolutionary selection pressures are congruent with the relationships identified by DRUMBEAT.

**Fig. 4.**
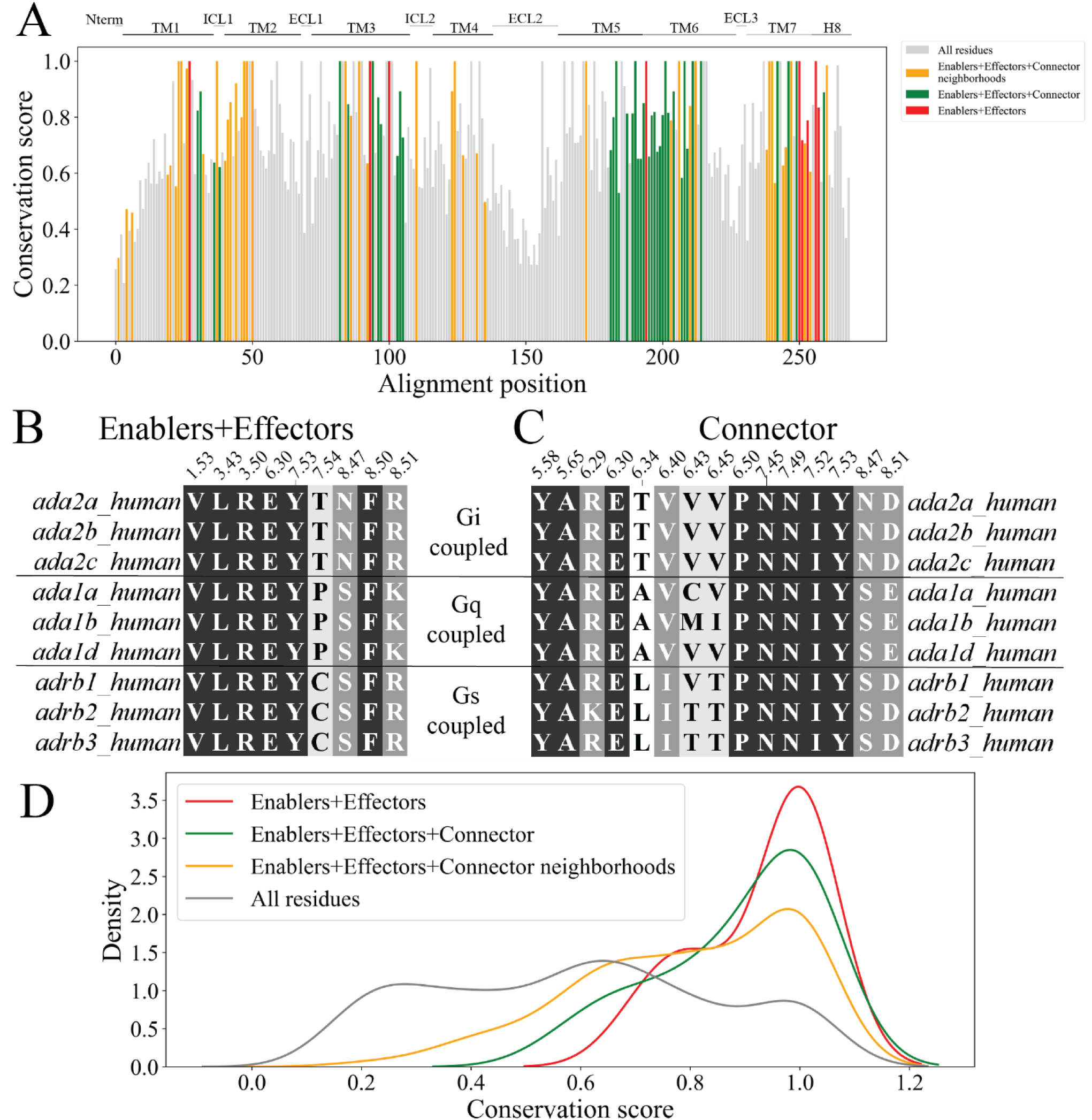
Evolutionary analysis of residue communities involved in conformational transitions in β_2_AR. (**A**) Mapping of the conservation score for all residues in human adrenergic receptors alignment. Sets of residues identified via DRUMBEAT are highlighted (yellow, green, and red) and show higher overall conservation score compared to all residues (grey). (**B**) Alignment of residues contained in the enabler and effector central contacts across human adrenergic receptors. (**C**) Alignment of expanded set of residues which contains the connector contacts between enabler and effector subnetworks. (**D**) Probability density function of conservation score computed using Von Neumann entropy for enabler+effector central contact residues (red), connector residues (green), entire neighborhood residues (yellow), and all protein residues (grey).

### Do the transition enabler and effector residues differ among Dopamine Receptor D_2_R and D_3_R subtypes?

Designing subtype-selective modulators and/or ligands targeting highly homologous proteins is one of the foremost challenges in the GPCR drug discovery. We applied DRUMBEAT to identify the residue communities involved in the deactivation transitions of D_2_R versus D_3_R, two prominent drug targets that so far have not been tractable to isoform-selective modulation.(*49*, *50*) Such an example would also demonstrate the broad applicability of the DRUMBEAT framework. We performed deactivation MD simulations on D_2_R and D_3_R generating 11 and 12 trajectory ensembles, respectively, with details provided in the Methods section. Preprocessing of the trajectory data included contact mapping and feature selection similar to those described for β_2_-adrenergic receptor, followed by constructing the BNM universal graph. We applied DRUMBEAT and identified relevant TRACs for each system.

We find that β_2_AR, D_2_R, and D_3_R exhibit both shared and distinct deactivation mechanisms. Firstly, we found that in the MD simulations, the deactivation of D_2_R is faster in time scale by an order of magnitude when compared to β_2_AR and D_3_R (0.1 μs vs 1-5 μs). Although the reported dopamine receptor MD simulations are shorter than β_2_AR, we extended the simulation time to ensure the receptor did not go back to intermediate or active states.

The top scoring residue contacts involved in D_2_R deactivation were identified using DRUMBEAT, with the subnetwork shown in Figure 5A. Five key contacts (TRACs) arise that span the intracellular and transmembrane regions of the receptor. The D131^3.49^_T68^2.38^ contact at the intracellular region is determined to be a key contact in both D_2_R and D_3_R. In the transmembrane region, D80^2.50^, S121^3.39^ and N422^7.49^ residues form two key contacts in D_2_R that show a hydrogen bond. D131^3.49^_T68^2.38^ contact plays an important role in D_3_R deactivation as well, along with 4 other contacts (subnetwork shown in Figure 5B). Interestingly, the contacts highlighted for the D_3_R system span the entire protein (See Figure 5E). Specifically, the contact F170^4.62^_F172^ECL2^ at the top of TM4 and in the ECL2 stands out in the D_3_R system. This contact is unique to the D_3_R and has been shown via mutation studies to play a significant role in ligand binding although it is not located directly in the ligand binding site.(*51*) The protein-spanning nature of TRACs in D_3_R alludes to the allosteric nature of the receptor’s activity. This is further corroborated by the fact that the neighboring contacts in the F170^4.62^_F172^ECL2^ TRAC are in both the extracellular and intracellular regions of the receptor (See Figure 8 in Discussion). The difference in sites of the D_2_R and D_3_R receptor TRACs manifests into distinct mechanisms by which these receptors deactivate.

**Fig. 5.**
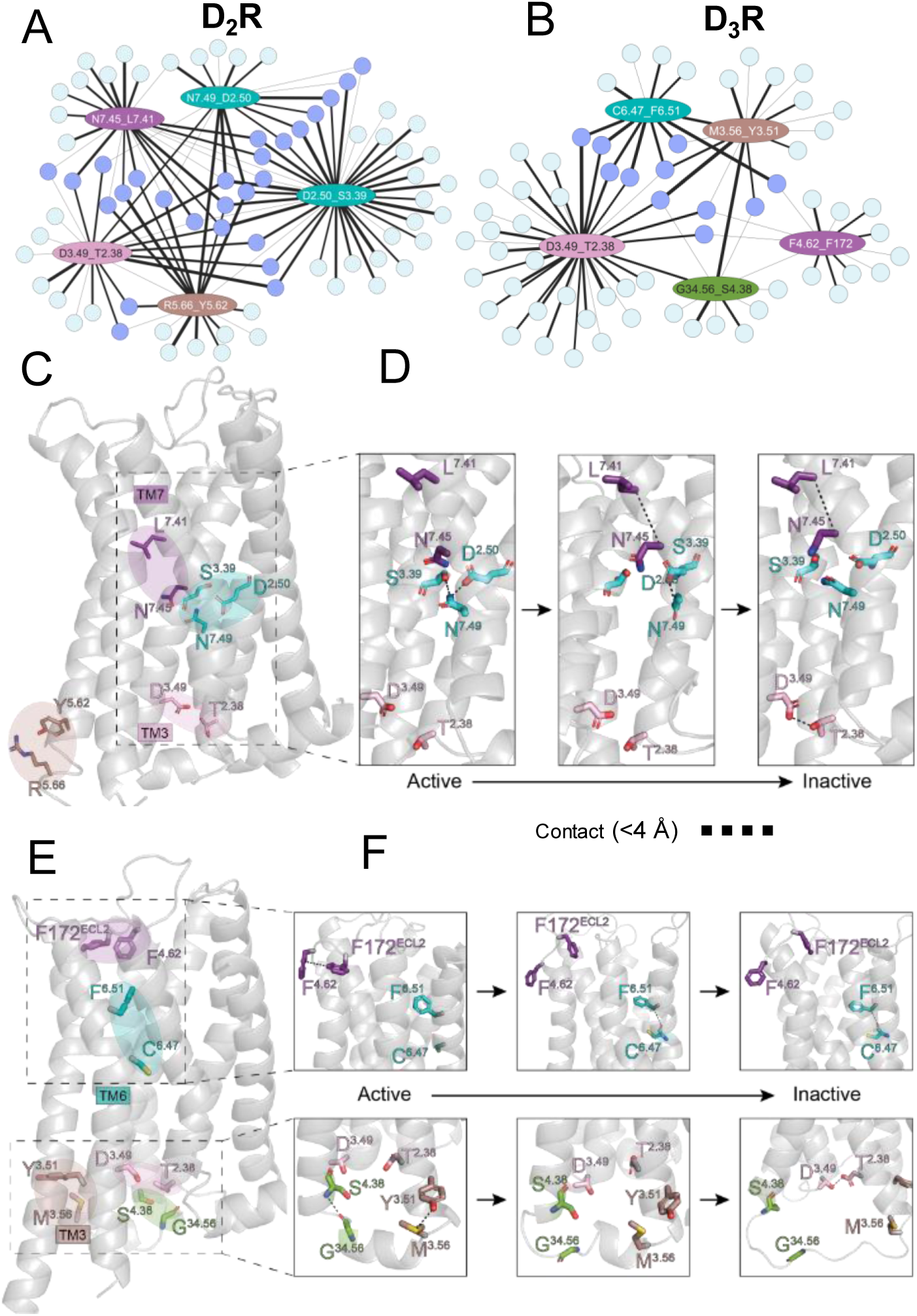
Application of DRUMBEAT to Dopamine D_2_ and D_3_ receptors. (**A** & **B**) Subnetwork from the universal graph for the top five ranked contacts for D_2_R (**A**) and D_3_R (**B**). Connector contacts shared between top ranked contacts are highlighted in purple. (**C**) Projection of top contacts on D_2_R structure. (**D**) Mapping of the structural changes that occur in D_2_R via changes in key contacts as identified by DRUMBEAT. (**E**) Structure of D_3_R with key contacts highlighted. (**F**) Mechanism of deactivation for D_3_R via changes in key contacts. Notably, contacts in both intracellular and extracellular regions initiate the transition for D_3_R. Both receptors transition into the inactive state via the formation of the D^3.49^_T^2.38^contact (shown in orange).

Even though D_2_R and D_3_R are highly similar in sequence and structure, DRUMBEAT uncovered distinct mechanisms by which each receptor transitions between active and inactive states. In D_2_R, initially the contacts between D80^2.50^, S121^3.39^, and N422^7.49^ residues stabilize the active state via hydrogen bonding. The first step in deactivation is mediated by formation of the N418^7.45^_L414^7.41^ contact at the kink region of TM7 (shown in red in Figure 5C and 5D). This most likely corresponds to the outward movement of TM7 that we observed in the β_2_AR study and is a preliminary step for deactivation. Then, hydrogen bonding between D131^3.49^_T68^2.38^ forms in the inactive state, possibly stabilizing the conformation. Interestingly, this contact also plays a similar role in D_3_R upon reaching the inactive state (See Figure 5F). The difference in the D_3_R deactivation mechanism arises from the other TRACs identified as central contacts.

Notably, initial transition out of the active state involves breaking of contacts in both the intracellular and extracellular region of the receptor, suggesting allosteric coordination (Figure 5F). The contact C341^6.47^_F345^6.51^ at the kink region of TM6 facilitates the final transition into the inactive state, corresponding to the helical movement of TM6 towards TM3, also in line with what we saw for β_2_AR as well as what is commonly reported for GPCR deactivation. Although the stability of the inactive state in both D_2_R and D_3_R is facilitated by the D131^3.49^_T68^2.38^contact, the active state is held together by a notably different set of residues and contacts. These findings point to evolutionary adaptations that likely enable functional specificity, despite overall receptor structure homology.

One final yet notable difference between β_2_AR and dopamine receptors is the behavior of the salt bridge formed between arginine on TM3 (R^3.50^) and glutamate on TM6 (E^6.30^). In β_2_AR deactivation trajectories, the salt bridge is consistently open during the active state and forms upon deactivation to the inactive state. Once in the inactive state, the salt bridge remains predominantly formed, with occasional fluctuations between open and closed states. These dynamics led to the contact being identified as a key TRAC by DRUMBEAT (R131^3.50^_E268^6.30^), as discussed above. In contrast, the dopamine receptor deactivation trajectories in this study reveal that the salt bridge forms in only a subset of trajectories (4 out of 11 for D_2_R; 5 out of 12 for D_3_R) and at a significantly lower contact frequency. This could imply that the salt bridge is more open in the inactive state(s) of dopamine receptors. To further investigate, we examined crystal structures of D_2_R and D_3_R in the inactive state (D_2_R: 6LUQ, 6CM4, 7DFP; D_3_R: 3PBL) for this salt bridge. We also analyzed the MD simulation trajectories of the antagonist bound inactive states of D_2_R and D_3_R. The crystal structures show the salt bridge is formed in two of three D_2_R inactive states (Fig. S6A) as well as in the D_3_R inactive state (Fig. S6B). In the inactive state MD trajectories, the salt bridge forms in approximately 12% of frames for D_2_R and 68% of frames for D_3_R (Fig. S6C). In the case of D_2_R, inactive state MD trajectories (initiated from the crystal structure – D_2_R: 6LUQ) show the salt bridge is formed initially, but then opens and remains so for the remainder of the trajectory. These findings indicate that while the salt bridge can form in the inactive state due to the proximity of TM3 and TM6, it is not essential for maintaining a stable inactive conformation. This suggests the existence of two distinct inactive states and a dynamic salt bridge between TM3 and TM6.(*52*, *53*)

### D_3_R transition is driven by subtype specific non-conserved residues

The findings from D_2_R and D_3_R suggest that while the dopamine receptors share some commonality in deactivation mechanisms, their variability could imply subtype-specific deactivation pathways that can be tapped into for subtype selectivity. To investigate this via evolutionary considerations, we divided residue positions into two sets: the set of "fully conserved" (C) positions and the set of "perfectly specific" (S) positions for a particular dopamine receptor class (e.g., D_2_R, D_3_R). As shown in the schematic in Figure 6A, the "fully conserved" positions remain invariant across all classes, whereas the "perfectly specific" positions uniquely distinguish one class of dopamine receptors from all the others. Purifying and adaptive selection are two prominent mechanisms shaping proteins’ evolution. Purifying selection removes harmful mutations from the gene pool and preserves specific residues in crucial positions within the protein, thereby maintaining key functionalities in protein structures. Adaptive selection, on the other hand, favors the fixation of advantageous mutations in the population. This process can lead to the neo-functionalization of paralogous genes, where new functions evolve. Purifying selection is likely the dominant evolutionary force acting on the "fully conserved" positions, as any alteration would be strongly selected against due to its impact on the core functionality of the receptor. In contrast, adaptive selection likely served as the primary evolutionary force acting on the "perfectly specific" positions, favoring those that provide a unique function to a given dopamine receptor class. Therefore, the ratio ρ=log_2_|S|/|C| provides an insight into the dominant evolutionary force acting on the residue community as identified by the DRUMBEAT analysis (i.e. key residues in the process of deactivation). Specifically, negative values denote receptors with mostly fully conserved positions and positive values denote receptors with mostly perfectly specific positions.

**Fig. 6.**
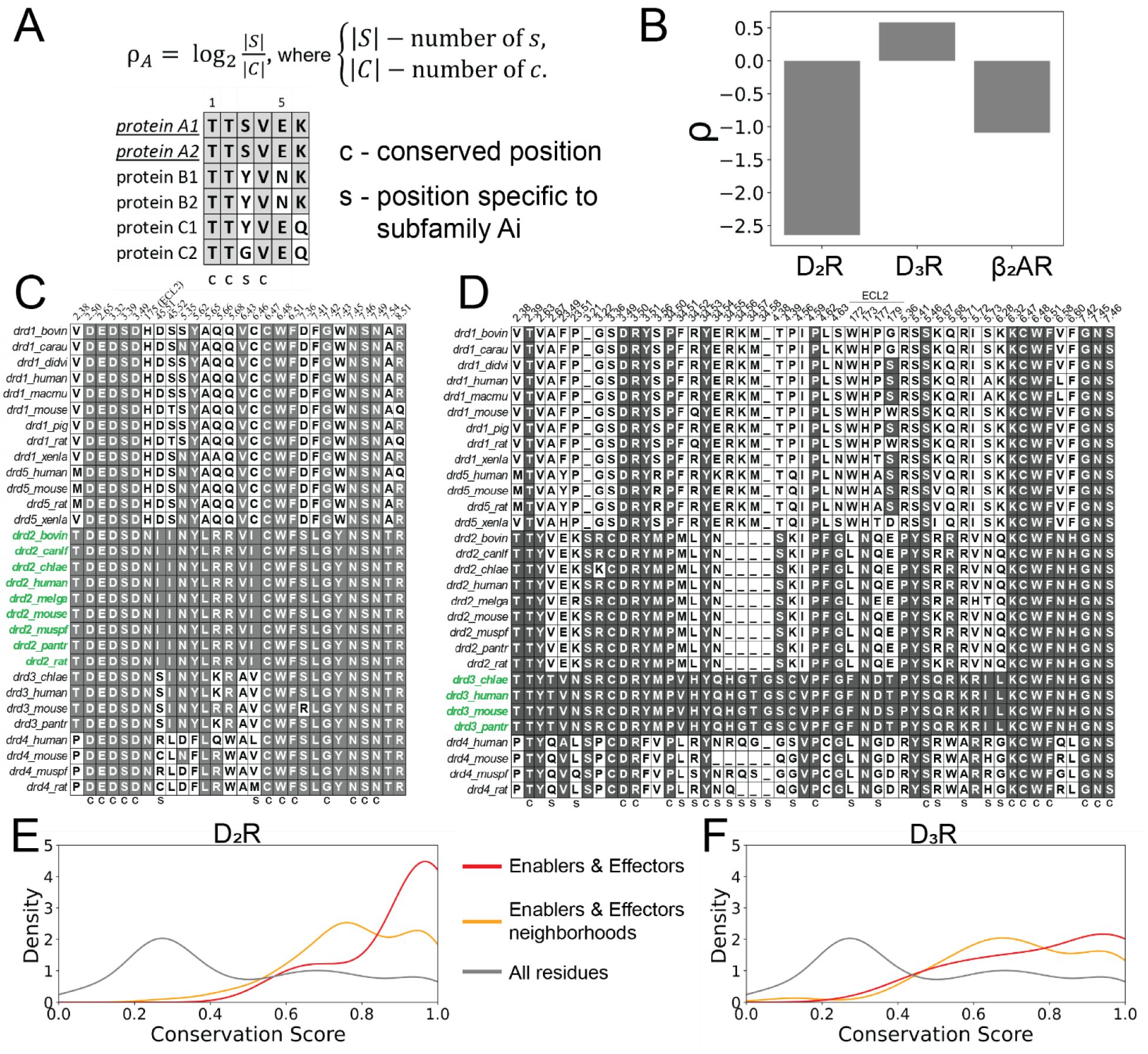
Evolutionary analysis of Dopamine receptors in accordance with findings from DRUMBEAT analysis. (**A**) Quantification of conservation and specificity fore residues. (**B**) Computed value for the ratio ρ for D_2_R, D_3_R, and β_2_AR. (**C** & **D**) Sequence alignment for D_2_R and D_3_R. (**E** & **F**) Conservation score for central contacts (red), central contacts and corresponding neighborhoods (orange), and all residues (grey).

To evaluate the conservation level of TRAC residues in dopamine receptors, we obtained MSA for all dopamine receptors (D1, D2, D3, D4, & D5) in all species from the GPCR database (www.gpcrdb.org). TRACs with a maximum weighted degree above 2.0 bits were extracted from each trajectory in the ensemble and the alignment was done using the residues in the union set of contacts. Alignment of residues from D_2_R TRACs is shown in Figure 6C, with D_2_R sequences highlighted in green. Similarly, Figure 6D shows the alignment for D_3_R TRAC residues, with D_3_R sequences likewise highlighted in green. For positions shown in Figure 6C and Figure 6D we calculated ρ and compared between receptors.

We computed ρ for all receptors in this study to provide a comparison of specificity (Figure 6B). We found that B_2_AR (ρ = −1.0) and D_2_R (ρ = −2.5) both exhibit negative ρ values, indicating that the key residues for deactivation are strongly conserved across the receptor class. In contrast, the key residues for D_3_R (ρ = 0.5) activation show a high degree of specificity, with nearly an order of magnitude difference, suggesting that these residues are evolutionarily diverging. Sequence conservation scores for D_2_R and D_3_R (Figures 6E and 6F) further reveal that while central contacts in D_2_R are highly enriched in conservation, those in D_3_R display only a modest increase, underscoring the subtype-specific nature of the D_3_R residues. To summarize, our analysis reveals that the key residues involved in D_3_R activation are characterized by positions that are evolutionarily diverging within the dopamine receptor family. Critically, this suggests that during evolution D_3_R began to diverge from its otherwise similar D_2_R counterpart by developing a unique activation pathway. Our DRUMBEAT-enabled ability to functionally characterize residues at these divergent positions provides direct insight into the functional mechanisms and evolutionary forces driving sub-functionalization, potentially inspiring novel drug molecule designs.

## Discussion

One of the major challenges in pharmacology is identifying target-specific drugs, especially for highly homologous proteins. To achieve target specificity, GPCR drug discovery paradigms in the last decade have pivoted to allosteric modulators.(*54*) However, understanding the functional impact of allosteric modulators on GPCR signaling is inherently limited by a dynamic nature of GPCR activation. Further, for multiple high-value drug target receptor families, including incretin, dopamine, serotonin, and muscarinics, isoform-specific residues are dispersed across the receptor structure complicating traditional structure-based strategies to realize isoform selectivity of small molecule therapeutics. In this study, we use a novel interpretable machine learning technique (Dynamically Resolved Universal Model for BayEsiAn network Tracking or DRUMBEAT) to identify spatially disperse, non-conserved residue communities that coordinate key conformational transitions in GPCRs. The distinct transition dynamics manifested by D_2_R and D_3_R provide a structural framework to identify selective allosteric sites and modulators for mitigating the opioid crises (D_3_R) and neuropsychiatric disorders (D_2_R).(*50*)

This work addresses a critical challenge in protein dynamics: identifying key regions and residues, or "allosteric residue communities”, that facilitate transitions between conformational states and ascertaining how these communities behave temporally. The DRUMBEAT methodology developed in this study uncovers the temporal dynamics and mechanistic underpinnings of conformational transitions, in the β_2_AR and two highly homologous and sought after drug targets, namely dopamine D_2_R and D_3_R receptors. We identified Time-Resolved Allosteric Communities (TRACs) that correspond to key sub-regions of the protein effecting the conformational changes and how their manifestations coincide with the transition of the receptor from active to inactive or vice versa. The results emphasize how probabilistic connectivities can identify critical contacts and their allosteric coordination during key transition events, providing a fresh perspective on receptor transition.

Two dominant computational approaches for studying protein dynamics are dynamic network analysis(*55*) and deep learning models.(*11*, *56*, *57*). Dynamic network analysis provides interpretable insights by constructing networks where nodes represent residues and edges are indicative of physical proximity. However, this method is inherently local, limiting its capacity to capture long-range allosteric interactions, especially those spanning spatially distant but functionally correlated regions. Conversely, deep learning methods offer high predictive performance(*11*) and scalability(*56*) but often lack interpretability as they provide, at most, feature importance rankings without mechanistic or interaction insights. Additionally, most of the deep learning models tend to rely heavily on extensive prior data, likely requiring thousands of MD trajectories in our case, and implying transfer learning with possible irrelevant false positives and hallucinations. The DRUMBEAT approach introduced in this study bridges these gaps, combining the interpretability of DNA with the scalability and data-driven nature of DL while offering a novel capability: time-resolved mechanistic analysis. A standout feature of this work is the temporal resolution afforded by DRUMBEAT, which reveals not just the key contacts and their involvement in transition but also the granular temporal anchoring and order of events during transitions. The temporal resolution allows for identifying enabler and effector residue communities, key subregions of the protein involved in the deactivation process.

Using DRUMBEAT to analyze the deactivation MD simulation trajectories of β_2_AR, we uncovered a mechanism of deactivation (Fig. 7A) where the TM7 is pulled out of the G-protein binding site by helix 8 through a transition enabler residue community around the contact C327^7.54^_F332^8.50^; this precedes the inward movement of TM6 movement towards TM3 effector community around R131^3.50^_E268^6.30^, a well-known indicator of the inactive state. This sequential relationship suggests a previously unknown causal mechanism, with the enabler’s coordinated motions serving as a precursor to the effector’s larger conformational changes.

**Fig. 7.**
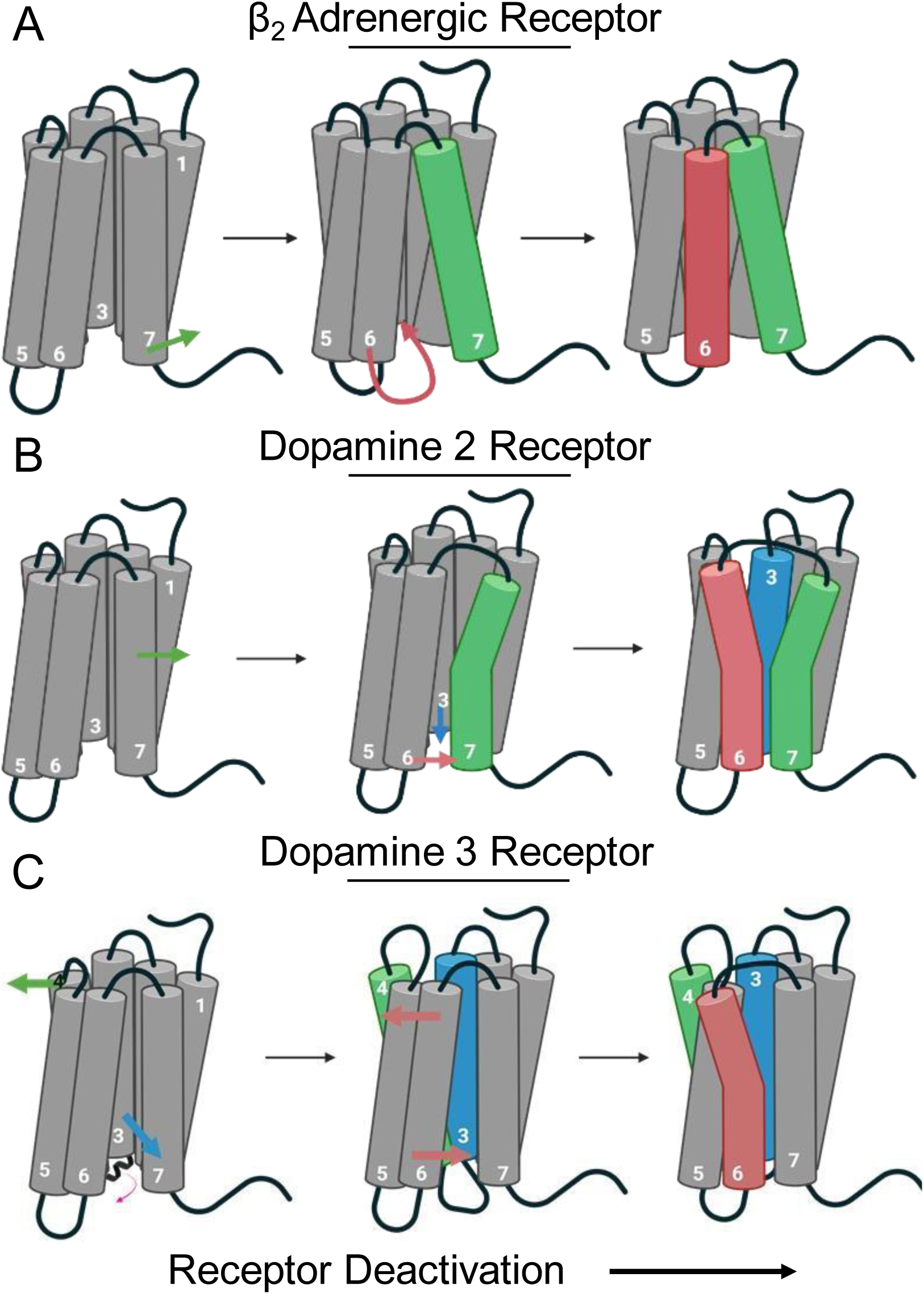
Schematics of the deactivation mechanisms identified by DRUMBEAT. (**A**) Deactivation mechanism for β_2_AR characterized by outward movement of TM7 followed by swinging of TM6 towards TM3. (**B**) D_2_R deactivation initiates via movement of TM7 at the central region of the receptor followed by TM3-TM6 closing. (**C**) D3R deactivation starts with allosteric movement of extracellular region of TM4 and intracellular region of TM3 and concludes with movement of TM6.

The DRUMBEAT analysis revealed further similarities and differences between the two dopamine receptors, D_2_R and D_3_R. We found that in D_2_R, the main mechanism of deactivation initiates at the central region of the receptor with movement of TM7 followed by TM3 and TM6 moving closer together (Fig. 7B). In contrast, the primary deactivation mechanism in D_3_R is characterized by an initial allosteric movement of the extracellular region of TM4 and intracellular region of TM3 which enables the movement of TM6 towards TM3 (Fig. 7C).

Despite these differences, these receptors share a fundamental reliance on specific contact networks for transitioning between states, illustrating both conserved and subtype-specific features in GPCR deactivation. The distinct deactivation pathway in the D_3_R receptor, together with strong enrichment in the “perfectly specific” positions, lead us to hypothesize that D_3_R receptors have significantly and concertedly diverged from other dopamine receptors, suggesting a process of neo-functionalization in the evolution of this receptor subtype.

Analysis of the differences in the TRACS between highly homologous D_2_R and D_3_R can be harnessed for design of subtype-specific ligands. As depicted in Fig. 8A, the residue contact F170^4.62^_ F172^ECL2^ shows long range dependencies on D127^3.49^_ Y138^34.53^ and S146^4.38^ _ G141^34.56^ TRACS to enable and effect transition to the inactive state. The TRAC S146^4.38^ _ G141^34.56^ is involved in the helix to coil transition of the intracellular loop 2 observed during deactivation(*25*). This underscores allosteric relationships during transition, such as the information flow between intracellular and extracellular regions that is specific to D_3_R and not D_2_R. Identification of putative small molecule binding sites using the FindBindSite program(*58*) (see Methods) shows (Fig. 8B) that there is a cryptic binding pocket formed near the residues that enable the deactivation transition of D_3_R. This is a proof of concept for using DRUMBEAT to identify the dynamic elements that differentiate two highly homologous receptors.

**Fig. 8.**
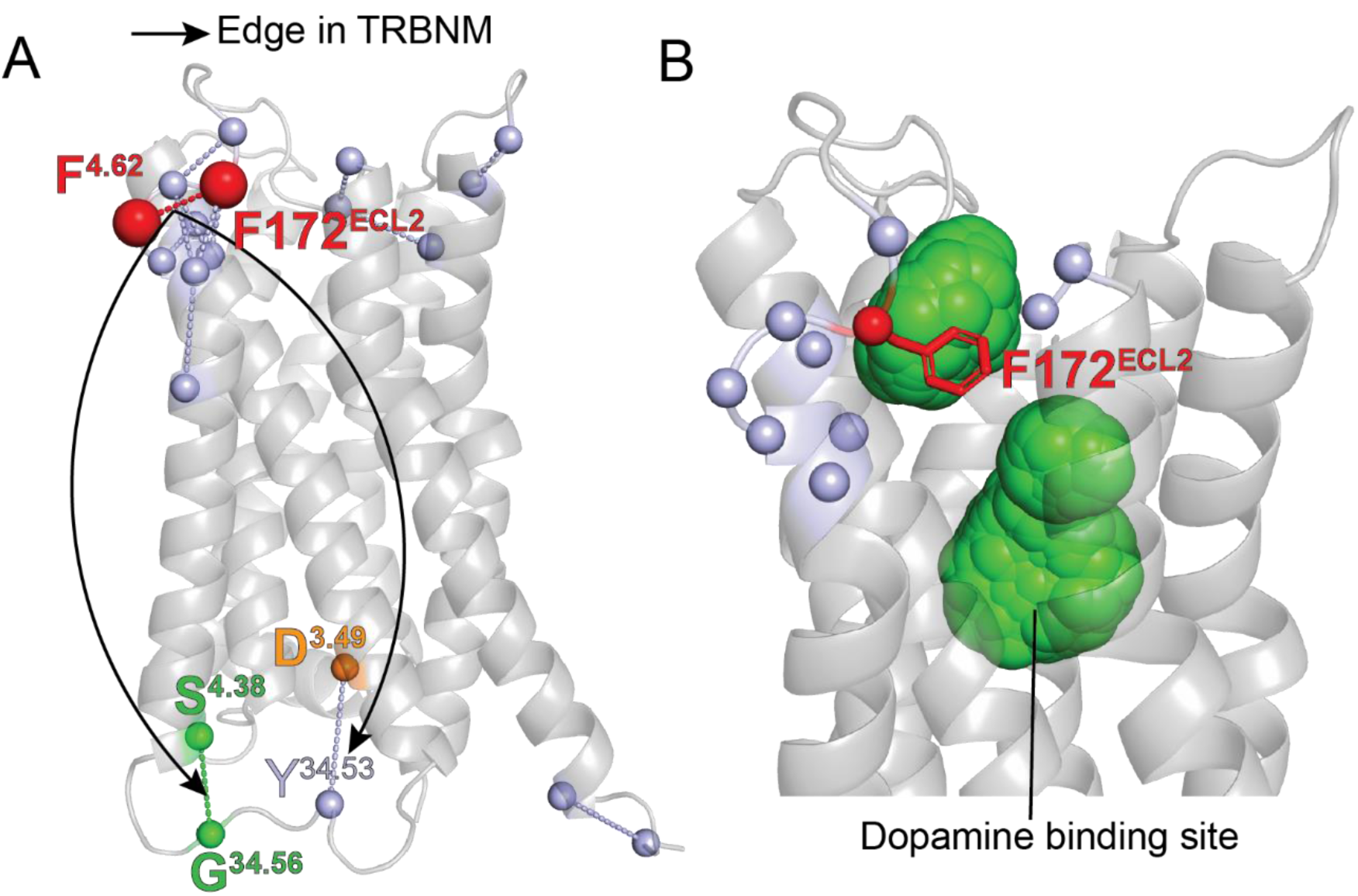
Putative D_3_R specific cryptic binding sites. (**A**) Residue contacts in the F170^4.62^_F172^ECL2^ TRAC shows direct allosteric co-dependencies on the intracellular TRACs specific to D_3_R identified in the DRUMBEAT analysis. (**B**) Green spheres are putative small molecule binding site identified using FindBindSite program.

Incorporating time steps into a BNM is generally done using dynamic Bayesian networks, which provide insight into the temporal relationships between the features of the system;(*59*) however, this approach is computationally extremely demanding and lacks precise time anchoring, making it suboptimal for our purpose of dissecting, at the granular level, the sequence of events that drive transitions. DRUMBEAT incorporates time in a notably different, but complimentary and equally mathematically rigorous, way that uses the BNM universal graph to identify all possible non-spurious dependencies and then precisely traces where the dependencies manifest alongside a time trajectory. Such granularity not only enhances understanding of GPCR dynamics but also lays the foundation for identifying intervention points in therapeutic contexts.

The implications of this work stem from both the innovative methodology and the results it produced. The DRUMBEAT approach introduces a novel way to study protein dynamics beyond their stationary states. By using a universal graph to establish probabilistic dependencies across conformational states and a scanning technique for temporal resolution, we identified key residues and events driving transitions. This dual capability provides critical insight into the most challenging regions of dynamic space—the transitions—by pinpointing when and where the protein deviates from equilibrium. Furthermore, the results propose new, collective reaction coordinates for transitions that move beyond traditional single-coordinate models (e.g., distances between two residues) to incorporate coordinated motions among multiple residues. This opens avenues for advanced sampling techniques to model transitions more efficiently. Lastly, the reproducibility of results across different GPCRs and the identification of conserved residues within evolutionary families underscore the methodology’s robustness. These findings not only shed light on the mechanics of GPCR transitions but also highlight residues critical for receptor family function, paving the way for experimental validation, mutation studies, and therapeutic targeting.

The DRUMBEAT approach presented in this work is a temporally resolved interpretable machine learning model designed to scale with expanding multimodal MD datasets, offering granular, mechanistic insights into protein allostery that have long been understudied. Although GPCRs—among the most complex allosteric proteins—serve as an ideal model system, the methodology is readily adaptable to other proteins and compounds, providing a novel, temporally resolved analysis that outperforms traditional network models and deep learning methods in its optimal balance of interpretability, precision, and efficiency. As data continue to grow, the linear computational complexity (given the universal graph) and transparent parametrization and hyperparametrisation of this approach ensure that its generalizability will only increase, paving the way for breakthroughs across a wide range of biological systems. Finally, the new algorithm is in principle applicable to any contiguous (temporal, spatial) and sufficiently granular data --- adaptation of DRUMBEAT to different such modalities will be a focus of our future studies.

## Acknowledgments

This work was funded by grants from the National Institutes of Health R01-GM117923 to NV, R01-LM013876 to NV, ASR, SBh, and SBr, and R01-LM013138 to ASR. The content is solely the responsibility of the authors and does not necessarily represent the official views of the National Institutes of Health. Additional support is acknowledged by Dr. Susumu Ohno Chair in Theoretical Biology (held by ASR), and Susumu Ohno Distinguished Investigator Fellowship (to GG).

## Funding

National Institutes of Health R01-GM117923 (NV)

National Institutes of Health R01-LM013876 (NV, ASR, SBh, and SBr)

National Institutes of Health R01-LM013138 (ASR)

Dr. Susumu Ohno Chair in Theoretical Biology (ASR)

Susumu Ohno Distinguished Investigator Fellowship (GG)

## Author contributions

Conceptualization: SS, NV, ASR, SBr

Data curation: BM, EM, GG

Formal analysis: BM, EM

Funding acquisition: SBh, NV, ASR, SBr

Investigation: BM, EM

Methodology: BM, EM, GG Software: BM, EM, GG

Supervision: SBh, NV, ASR, SBr Visualization: BM, EM

Writing – original draft: BM, EM, SBh, NV, ASR, SBr

Writing – review & editing: BM, EM, SBh, SS, NV, ASR, SBr

## Competing interests

Authors declare that they have no competing interests.

## Data and materials availability

All materials needed to evaluate the conclusions in the paper are outlined in the paper and/or the supplementary materials. All materials in this paper will be shared upon request. DRUMBEAT Code can be found on Github (https://github.com/bandyt-group/drumbeat). MD Simulation data for β_2_AR can be found at PNAS article (Ref 25; doi: 10.1073/pnas.1110499108)

## Supplementary Materials

Materials and Methods

Figs. S1 to S7

References (*60*-*69*)

## Notes

https://github.com/bandyt-group/drumbeat

### Competing Interest Statement

The authors have declared no competing interest.

