## Supplementary Materials for "Temporally Resolved and Interpretable Machine Learning Model of GPCR conformational transition"

**The PDF file includes:**

Materials and Methods
Figs. S1 to S7

### Materials and Methods

Preparation of Protein Structures for MD simulations

MD simulations were set up using experimental structures specific to each protein: $D_2R$ (PDB ID: 8IRS)(*26*) and $D_3R$ (PDB ID: 8IRT)(*26*). Agonists and G proteins were excluded from the structures. Missing side chains and loops, involving fewer than five residues, were reconstructed in the 3D models using Schrödinger's Maestro software (Schrödinger Release 2020-1, Schrödinger, LLC, New York, 2020). Local minimization was performed for residues within 5 Å of the mutation sites using MacroModel, while maintaining position restraints on all backbone atoms. Neutral acetyl and methylamide groups were used to cap protein termini, and histidine protonation states were determined using Maestro's protein preparation wizard. Receptors were inserted into an explicit POPC bilayer membrane using the PPM 2.0 functionality of the Orientation of Proteins in Membranes (OPM) tool.(*60*) The system was then solvated with TIP3P water and 0.15 M NaCl, prepared through CHARMM-GUI.(*61*, *62*) Final simulation boxes measured approximately 90 Å × 90 Å × 130 Å, and all proteins were modeled using the CHARMM36m force field.(*63*)

Details of the MD Simulations

MD simulations were performed using GROMACS 2022(*64*) with an integration timestep of 2 fs. The prepared systems underwent an initial energy minimization phase, applying position restraints of 10 kcal/mol·$Å^2$ to the heavy atoms of proteins, ligands, and lipids where applicable. This was followed by a 1-ns heating phase in the NVT ensemble, gradually increasing the temperature from 0 K to 310 K using the Nosé-Hoover thermostat. Subsequent equilibration in the NPT ensemble involved an initial 1-ns run with the same 10 kcal/mol·$Å^2$ restraints, which were progressively reduced (10, 5, and 1 kcal/mol·$Å^2$) over 5-ns intervals. A final 50-ns equilibration phase was conducted without restraints. The resulting configuration from this phase served as the starting structure for thirty-five production simulations, each initialized with distinct random velocities. Pressure was maintained at 1 bar using the Parrinello-Rahman(*65*) barostat, while nonbonded interactions were calculated with a 12 Å cutoff. Long-range electrostatics were treated using the Particle Mesh Ewald (PME) method,(*66*) and the LINCS algorithm was employed to constrain all bonds and water molecule angles.

MD data for $\beta_2$ Adrenergic Receptor

MD trajectories of deactivation of $\beta_2AR$ were procured from DE Shaw and Associates; these simulations were performed and published previously.(*25*) An ensemble of trajectories that showed transition from active to inactive was used for each system. We utilized 14 productive trajectories of condition A (protonated D130).(*25*)

Calculating Fingerprints of Inter-protein Pairwise Interactions

Pairwise residue contacts between protein residues were analyzed using the Python script library "GetContacts"(*67*) (https://www.github.com/getcontacts). This tool was employed to identify different types of interactions, including salt bridges (cutoff < 4.0 Å), hydrogen bonds (cutoff < 3.5 Å with an angle < 70°), van der Waals contacts (difference < 2 Å), π-stacking interactions (distance < 7.0 Å with an angle < 30° between aromatic planes), and cation-π interactions (distance < 6.0 Å with an angle < 60°). The analysis was conducted over the whole trajectory, excluding lipids, water and ions. Both atom selection groups were aligned with receptor residues. Custom Python scripts were used for one-hot encoding, generating binary contact fingerprints for each simulation frame, where "1" indicates a contact and "0" represents its absence.

Preprocessing of the MD simulation Data

Given $C^{all} = \{c_i\}$, the set of all contacts between two residues in a protein, the following steps were used for feature selection. First, contacts between neighboring residues were excluded from the data. Second, within each trajectory, contacts providing little to no information on other contacts were omitted. Specifically, pairwise mutual information was computed between contacts in each trajectory. Whichever contacts failed to have a pairwise mutual information value above 0.01 bits were also removed. This procedure is implemented in BaNDyT software.(*27*) Finally, the remaining features (contacts) were chosen in the previous steps across all individual trajectories to obtain the set of the most significant contacts $C = \{c_i\}$ with $C \subset C^{all}$.

The universal dataset $D^U = \{c_{ir}\}$ with $c_{ir}$ representing the state of the contact *i* ($c_{ir} \in \{0,1\}$) at each frame ($r$) is obtained by 1) concatenating all trajectories or 2) random sampling of frames from the individual trajectories. In general, the first technique is preferred to avoid data loss, but given long trajectories with many frames, concatenation results in a dataset that is too large for the BNM construction in a reasonable timeframe.

In our practice, given that the Bayesian network topology recovery is generally worst-case NP-hard, we found a dataset of roughly 1000 contacts and 15,000 frames to be near the limit of computational feasibility (one week on a HPCC). For all three receptors, the sampling technique was carried out to provide a single dataset for each system. Since we were interested in specifically studying the transition of the receptor, an enriched sampling was carried out around the transition point for each trajectory, to strengthen potentially borderline edges in the resulting BNM structures. Using the interhelical distance as a proxy for transition, states were randomly sampled (with replacement) from a region of 5000 frames centered at the drop in interhelical distance. The number of states sampled per trajectory was defined by the total number of trajectories for each system and the upper limit of 15,000 frames. For instance, 1200 frames were sampled from each of the 12 trajectories for $\beta_2$AR giving rise to a total of 14,400 frames. Similarly, 1400 frames were sampled from each $D_2$R trajectory, and 1200 frames were sampled from each $D_3$R trajectory, yielding 15,400 and 14,400 frame datasets, respectively. The resulting universal dataset served as an input for the BNM universal graph construction.

### Bayesian Network Model

For a given set of contacts $C$ and a universal dataset $D^U$ constructed as described above, we used BaNDyT software(*27*), a specialized implementation of BNomics(*68*, *69*) for MD simulations, to build the Bayesian network for each system. Let $\mathcal{G} = (C, \mathcal{E})$ be a Directed Acyclic Graph (DAG) model with $C$ the set of contact and $\mathcal{E} = \{e_{ij}\}$ the set of edges between contact *i* and *j* obtained using BaNDyT, then, the joint probability $\mathcal{P}(C)$ can be factorized as:

$$\mathcal{P}(C) = \prod_i p(c_i | pa_{c_i}),$$

with $pa_{c_i} \subseteq C$ the set of all parents of $c_i$ in $\mathcal{G}$. This equation states that the joint probability can be represented as the product of the conditional probabilities of each $c_i$ given its parents $pa_{c_i}$.

### Dynamically Resolved Universal Model for Bayesian Network Tracking

A Dynamically Resolved Universal Model for BayEsiAn network Tracking (DRUMBEAT) was built for *each* trajectory ($k$) using the universal graph, $\mathcal{G}$. Specifically, given MD simulation data $D^k$ of a trajectory and the universal graph, $\mathcal{G}$, DRUMBEAT uses a sliding window strategy to detect if a specific direct dependency is present in a given temporal interval

(2δ) from row in $D^k$. A time-dependent Mutual Information (MI) serves as a non-parametric measure to quantify the dependency strength between the two nodes connected by an edge in $\mathcal{G}$ within the temporal interval $\mathrm{t} - \delta$ to $\mathrm{t} + \delta$:

$$\mathrm{M}I_t(i,j|\delta) = \mathrm{H}\left(X_{i,t-\delta:t+\delta}\right) + \mathrm{H}\left(X_{j,t-\delta:t+\delta}\right) - \mathrm{H}\left(X_{i,j,t-\delta:t+\delta}\right)$$

An optimization procedure is applied for each edge to determine the optimal window size by maximizing MI across all possible window sizes. Multiple window sizes were used, ranging from 150 to 1050 timesteps.

Finally, we define the Time Resolved Allosteric Community (TRAC) for the contact *i* as the time resolved weighted degree given the universal graph $\mathcal{G}$:

$$\mathrm{TRAC}_i(t) = \sum_{j \in \Gamma_i} MI_t(i,j|\delta),$$

with $\Gamma_i$ being the neighbors of the vertex i in $\mathcal{G}$.

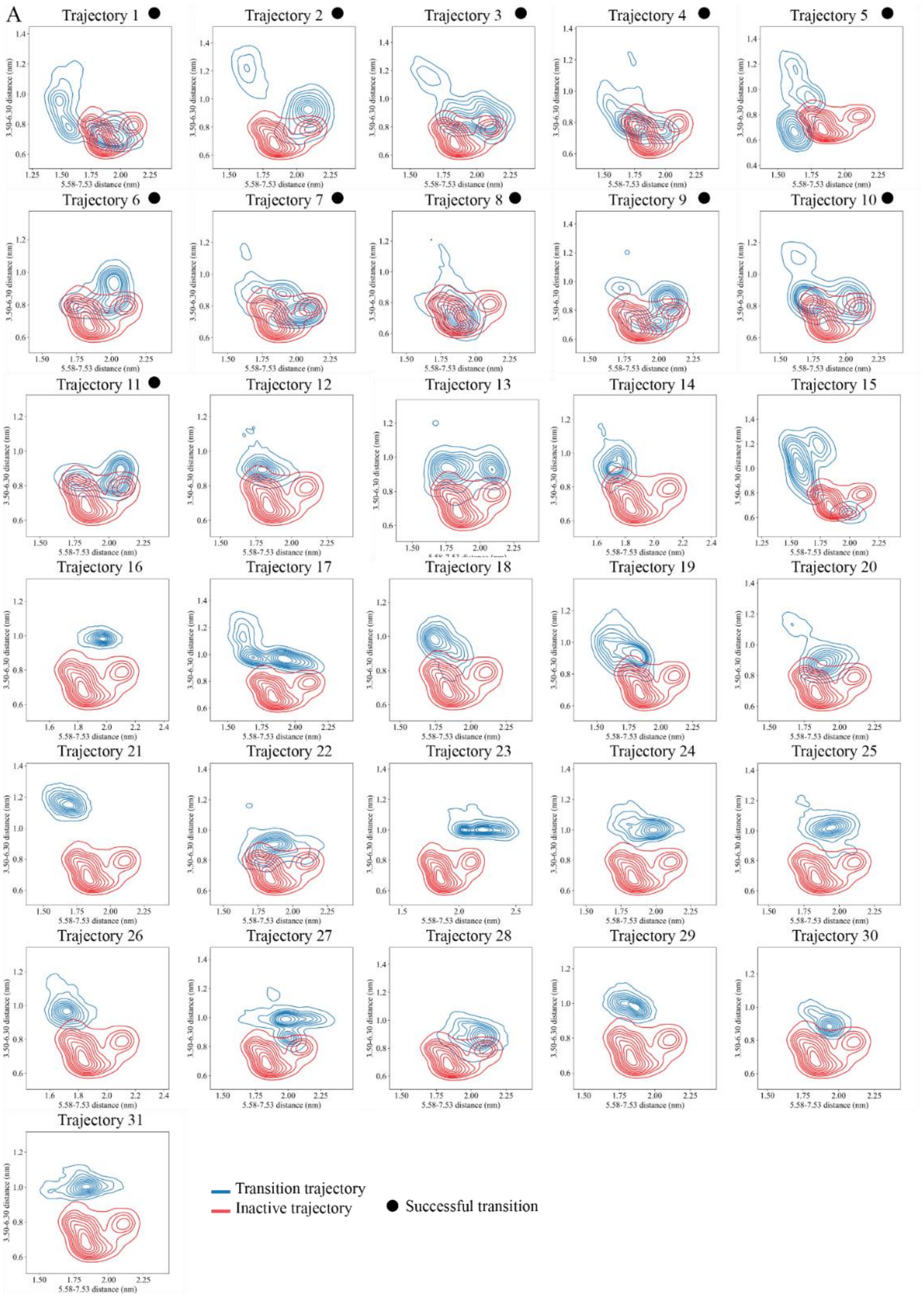


**Fig. S1.**

**Density distribution of inter-residue distances between TM5-TM7 and TM3-TM6 from 31 transition trajectories and inactive state trajectories of $D_2$DR.** Kernel density estimation plots show the relationship between the TM5-TM7 (5.58-7.53) and TM3-TM6 (3.50-6.34) distances measured in nm. The blue contours represent the density distribution of transition trajectories, while the red contours correspond to the inactive trajectory. Black dot - successful transition trajectory.

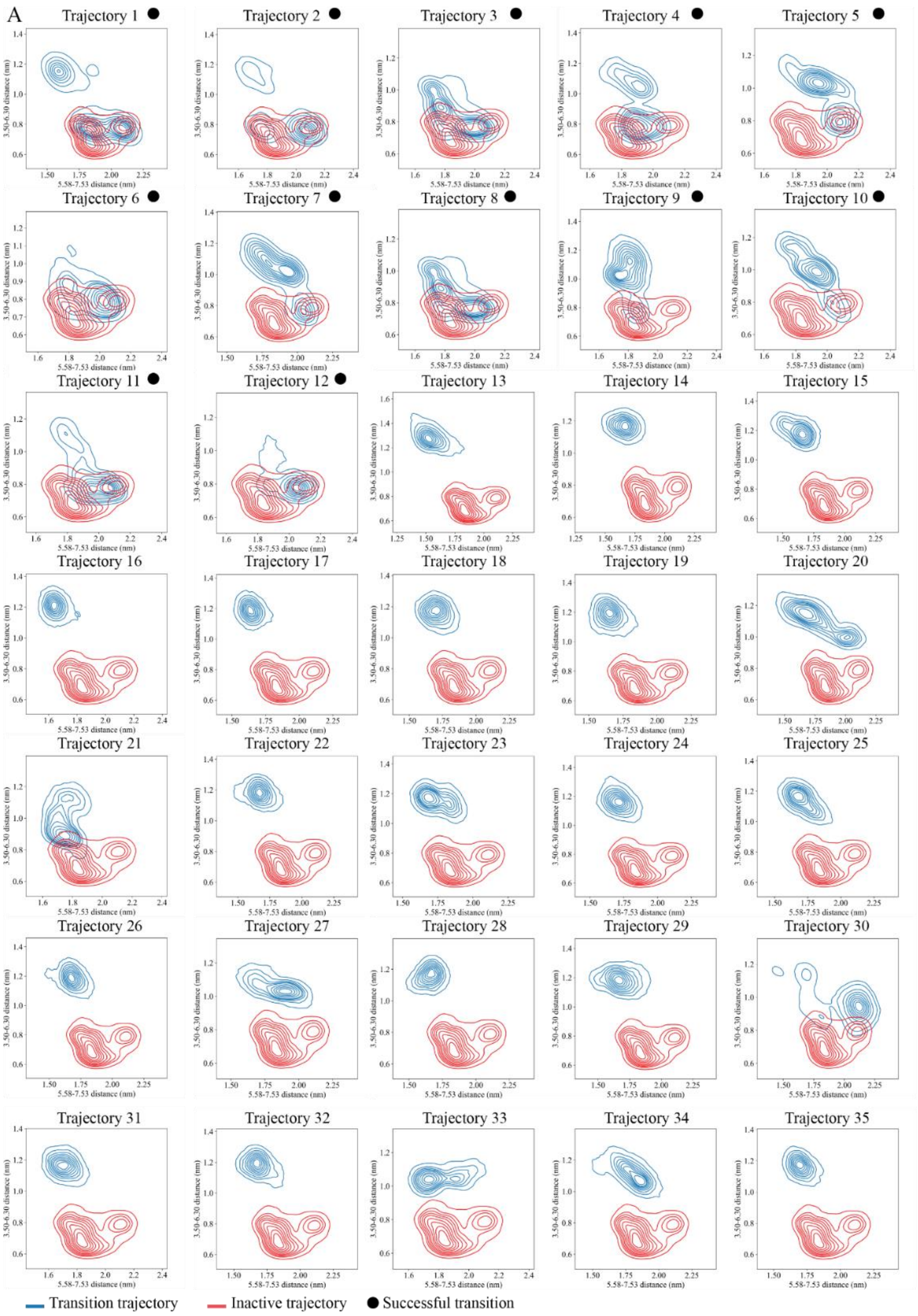


**Fig. S2.**

**Density distribution of inter-residue distances between TM5-TM7 and TM3-TM6 from 31 transition trajectories and inactive state trajectories of $D_3$DR.** Kernel density estimation plots show the relationship between the TM5-TM7 (5.58-7.53) and TM3-TM6 (3.50-6.34) distances measured in nm. The blue contours represent the density distribution of transition trajectories, while the red contours correspond to the inactive trajectory. Black dot - successful transition trajectory.

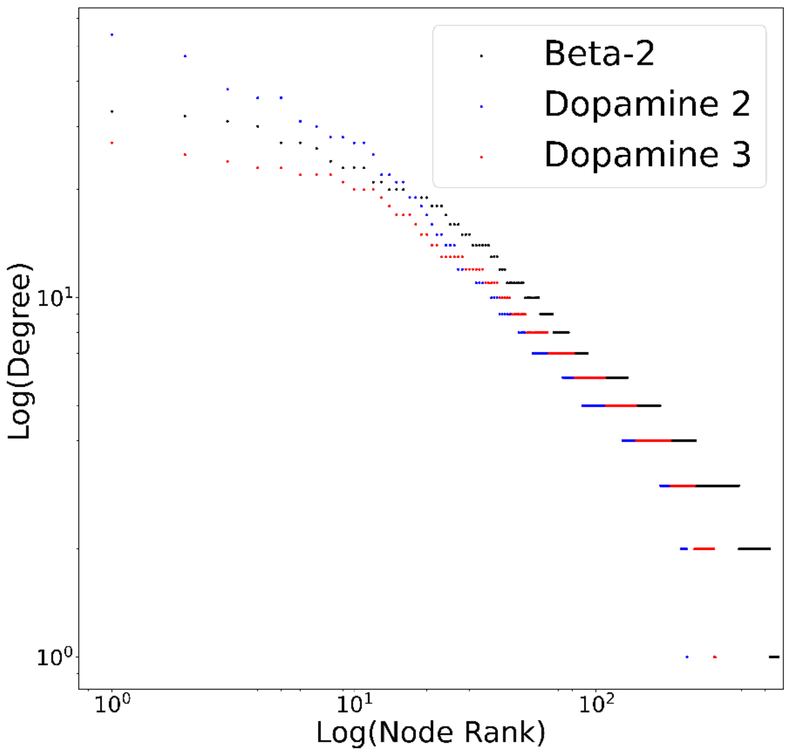


**Fig. S3.**

Degree distribution as scatter plot for all GPCR systems.

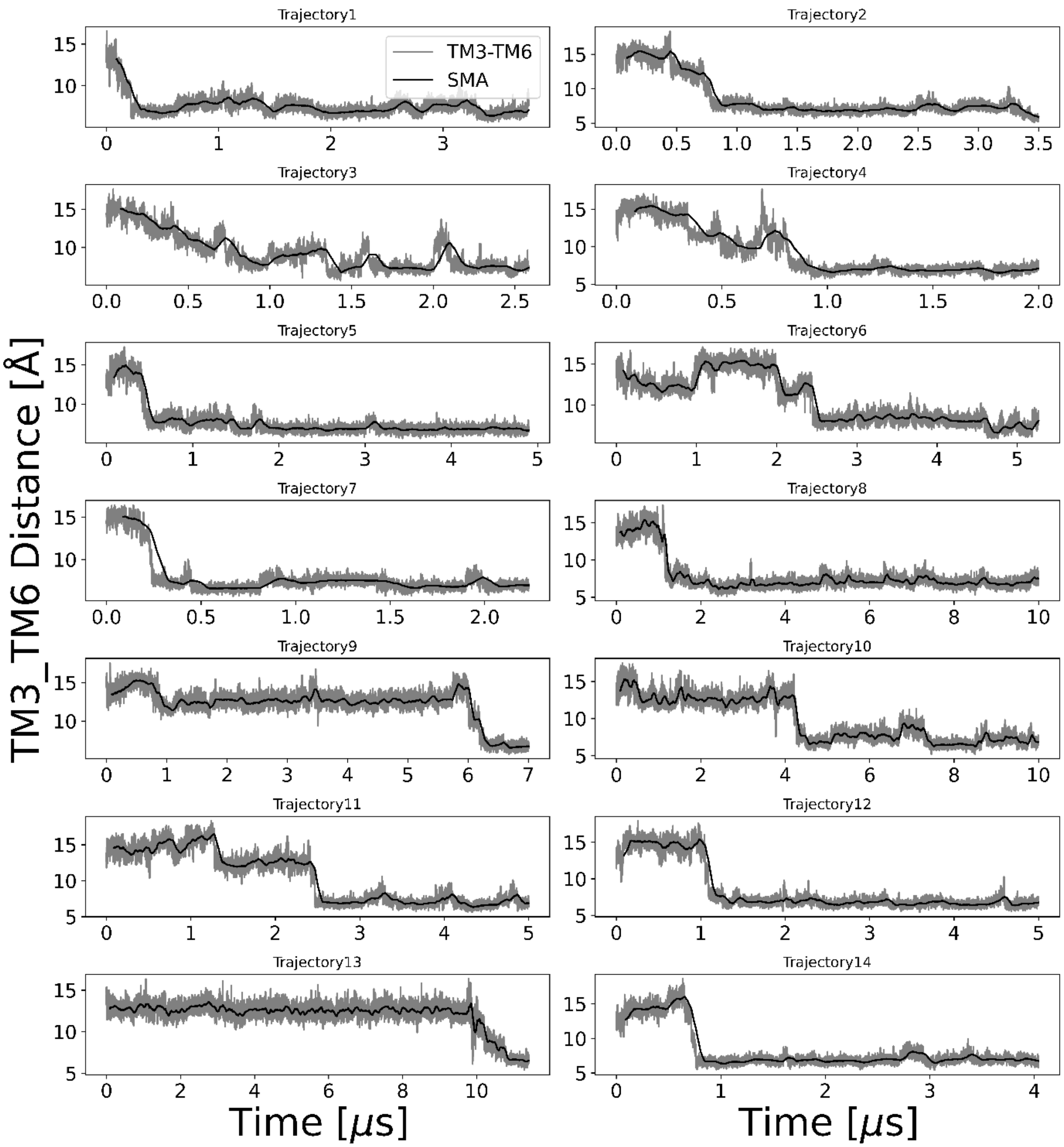


**Fig. S4.**

TM3-TM6 distances for all β2AR trajectories showing transition in all cases.

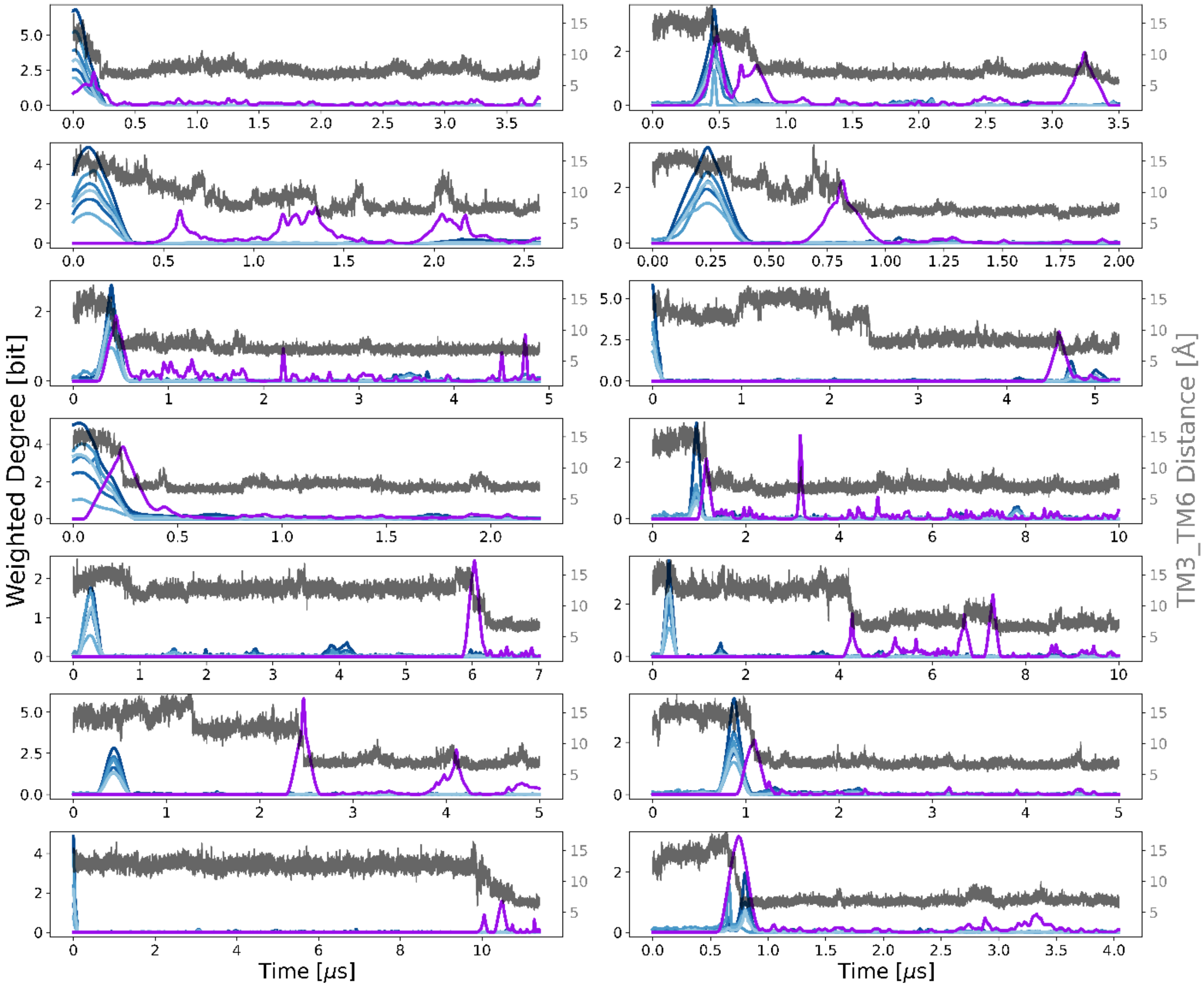


**Fig. S5.**

Weighted degree plots for all β2AR trajectories.

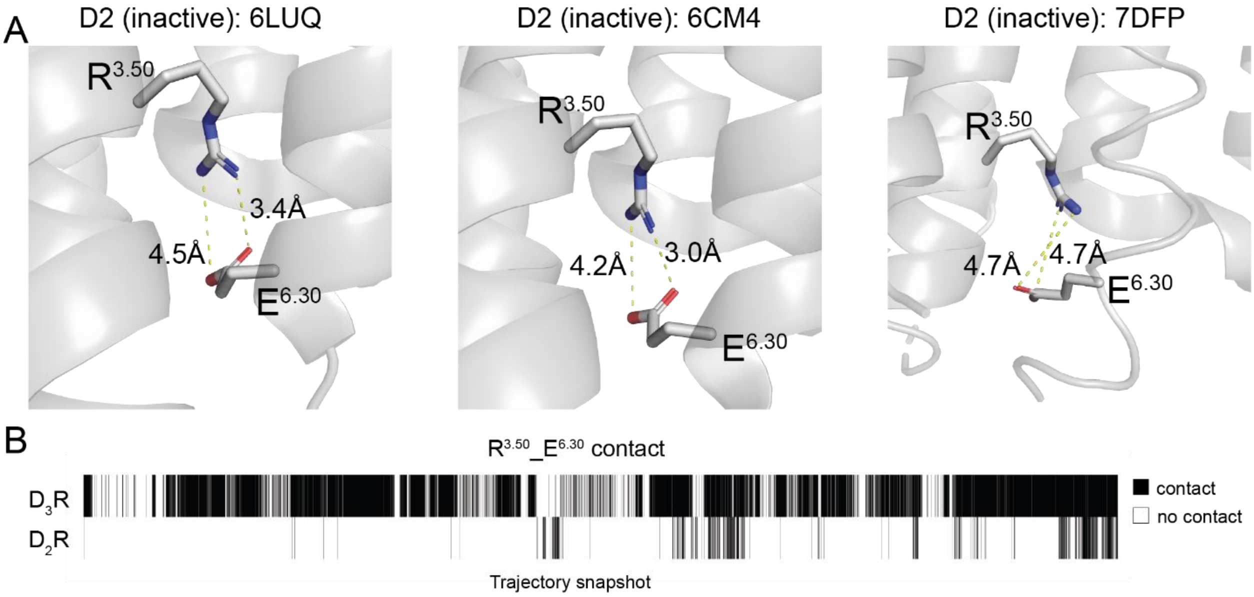


**Fig. S6.**

**A.** Structural representation of three available crystal structures of D2R demonstrating the minimal distances between R*3.50* and E*6.30* residues. **B.** Heatmap of R*3.50*_E*6.30* contact appearance throughout $D_2R$ and $D_3R$ inactive state ensemble trajectories. Black color indicates formation of contact in the trajectory snapshot, white – absence.

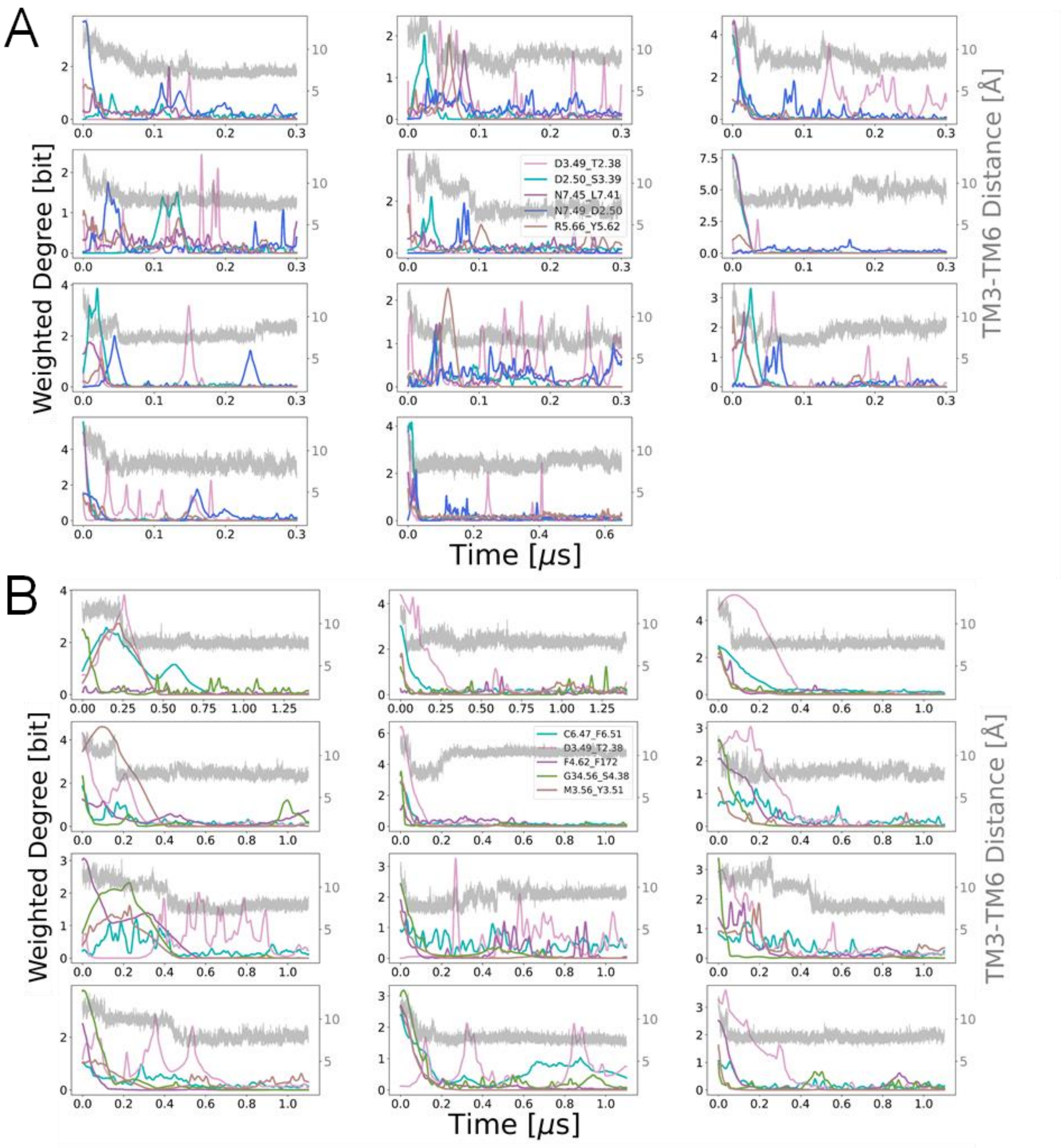


**Fig. S7.**

Weighted degree plots for all (A) $D_2R$ trajectories and (B) $D_3R$ trajectories. Top five ranked nodes are plotted for each system.
